# A *multi-track* landscape of haematopoiesis informed by cellular barcoding and agent-based modelling

**DOI:** 10.1101/2024.03.28.587126

**Authors:** Dawn S. Lin, Stephen Zhang, Jaring Schreuder, Jessica Tran, Toby Sargeant, Don Metcalf, Ashley P. Ng, Tom S. Weber, Shalin H. Naik

## Abstract

In classic ‘ball-and-stick’ models of haematopoiesis the implicit assumption is that all cells within each defined stem or progenitor cell population are equivalent in their fate. Instead, more recent models suggest a haematopoietic stem and progenitor cell (HSPC) ‘continuum’ of lineage bias and commitment, which is largely inferred through ‘snapshot’ analysis of single cell gene expression or clonal fate. However, the dynamic assessment of lineage commitment of specific HSPC populations and their clonal output over time *in vivo* is still lacking but is essential to fully inform accurate models of haematopoiesis. Here, using cellular barcoding we compare the single cell output of long-term haematopoietic stem cells (LT-HSCs), short-term HSCs (ST-HSCs), multipotent progenitors (LMPPs), common myeloid progenitors (CMPs), common lymphoid progenitors (CLPs), and macrophage/dendritic cell progenitors (MDPs). Each population was assessed for their output to multiple haematopoietic cell types spanning a subset of time points from 9 to 112 days of haematopoiesis after transplantation. These analyses revealed a wide range of clonal fate patterns that were inconsistent with their eponymous labels, i.e. stem and multipotent progenitors were rarely multi- or equipotent, and ‘common’ progenitors were often highly restricted in their fate. To better describe how these clonal patterns integrate into a revised landscape, a novel agent-based mathematical modelling approach that explicitly accounts for haematopoiesis at a clonal level was developed to allow the simulation of growth, timing and branching of clonal trajectories that underlie the process. Rather than a continuum, the proposed model is suggestive of multiple tracks down which clonal trajectories progress, and where fate can branch to a track of lower potency at multiple points down the entire cascade of haematopoiesis.

**Graphical abstract:** 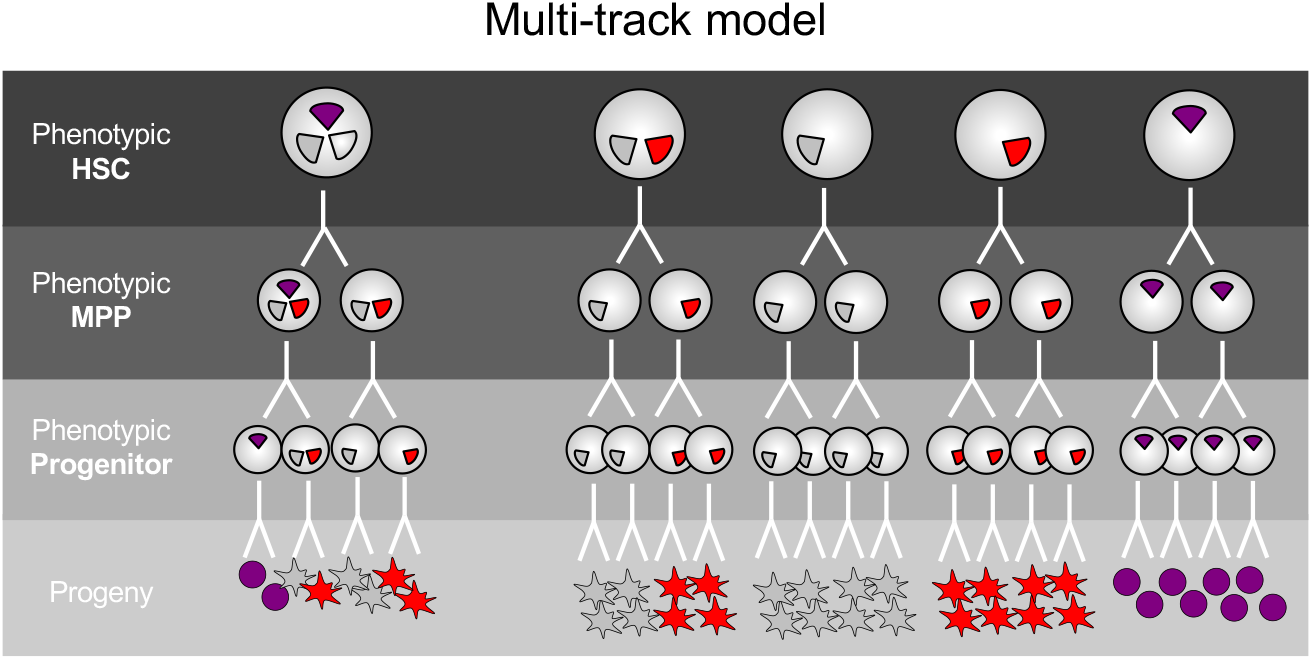

In a multi-track model, while some HSPCs are multipotent and branch into daughters with committed fate (left trajectory), other HSPCs may already committed or biased to a certain lineage such that their daughters inherit and maintain it in subsequent divisions during haematopoiesis. However, this commitment/fate bias is not evident through current phenotypic definitions of HSPC subsets (background colour) but occurs through putative expression of transcription factors, epigenetic programming or other factor that is currently unresolved (as indicated by the coloured fate potential triangles inside cells).

## Introduction

Generating red and a multitude of white blood cell types in balanced numbers relies on coordinated processes in haematopoiesis. An enduring model is that all cell types derive from multipotent self-renewing long-term haematopoietic stem cells (LT-HSCs) that reside at the apex of haematopoiesis (Orkin and Zon, 2008; Weissman and Shizuru, 2008). Multiple haematopoietic stem and progenitor cell (HSPC) populations exist downstream of LT-HSCs, with each successive HSPC stage progressively restricted in the number of cell types (potency) and number of progeny (cellular output) they generate.

Population-level analyses of potency and production has given rise to classical models that conceptualise HSPCs as ‘multipotent’ or ‘common’ precursors to their prescribed cell types. Hence the terminology multipotent progenitor (MPP) (Adolfsson et al., 2005; Cabezas-Wallscheid et al., 2014; Pietras et al., 2015; Wilson et al., 2008), common myeloid progenitor (CMP) (Akashi et al., 2000), common lymphoid progenitor (CLP) (Kondo et al., 1997), etc. Such assertions, however, had led to fixed ‘ball-and-stick’ models of haematopoietic relationships wherein the implicit assumption is that every cell within that population has similar potency and cellular output. However, while these models have served the community well, numerous studies now demonstrate that current immunophenotype-based definitions of HSPC subpopulations are inadequate in capturing fate heterogeneity, thereby rendering the ball-and-stick visualisations of haematopoiesis overly simplistic, and extensively reviewed in (Ema et al., 2014; Haas et al., 2018; Jacobsen and Nerlov, 2019; Laurenti and Göttgens, 2018).

Instead, the utilisation of i) single cell ‘omics (scRNA-seq, scATAC-seq) and ii) clonal lineage tracing have highlighted both molecular and fate complexity within HSPC populations, respectively. Single cell ‘omics approaches are attractive in that cells of similar transcriptional states can be grouped (leading to identification of novel HSPC subsets) (Amann-Zalcenstein et al., 2020; Giladi et al., 2018; Olsson et al., 2016; Paul et al., 2015; Triana et al., 2021; Zhang et al., 2024) or ordered (leading to inferred trajectories of haematopoiesis) (Buenrostro et al., 2018; Herman et al., 2018; Kucinski et al., 2024; Pellin et al., 2019b; Tusi et al., 2018) in an unbiased way. However, an important caveat is that a stem/progenitor’s transcriptional profile does not necessarily correlate with its function, *i.e.* the cell types it makes. In other words, what a cell transcriptionally *is* does not necessarily correlate with what it *does*. Indeed, studies pairing transcriptional states with fate at a clonal level using SIS-seq (Luyi et al., 2018; Tian et al., 2021) or similar approaches (Biddy et al., 2018; Emert et al., 2021; Rodriguez-Fraticelli et al., 2020; Weinreb et al., 2020) provided powerful new insights, including the surprising result that genes that predict fate bias or other clonal properties may not be readily identified through single cell ‘omics approaches in isolation. Therefore, clonal lineage tracing is the only way to ascertain the actual (not inferred) lineage fate of single HSPCs (Naik et al., 2014). This strategy has been used to functionally define HSPC fate heterogeneity in vitro (Karamitros et al., 2018; Lee et al., 2017; Lin et al., 2018; Notta et al., 2016; Velten et al., 2017) and in vivo (Lu et al., 2011; Naik et al., 2013; Pei et al., 2017; Rodriguez-Fraticelli et al., 2018; Yu et al., 2016).

Together, these studies have challenged the assumptions underlying ball-and-stick models of haematopoiesis and highlight that a clone-centric approach needs to be incorporated into contemporary models. Many such models have converged on the term ‘continuous’ or ‘continuum’ model proposed by Velten *et al*. where the authors elegantly demonstrated heterogeneity in HSPC potential at a clonal level and linked them *in silico* to transcriptomic landscapes (Velten et al., 2017). They proposed a ‘continuum of low-primed undifferentiated haematopoietic stem and progenitor cells’ or ‘CLOUD-HSPCs’. However, this terminology has been open to interpretation and “the degree to which early haematopoiesis is characterized by a continuum versus distinct populations therefore remains a question that requires further investigation” (Laurenti and Göttgens, 2018). For example, it is not clear whether there exists a *horizontal* continuum of low-primed undifferentiated or semi-stable HSPC clouds in which cells “gradually acquire continuous lineage priming that nudges them toward” lineage fate (Hirschi et al., 2017) or whether multiple *vertical* tracks of pre-committed/biased HSPCs with different fate patterns occur from which the entire haematopoietic pathway descends (Fig 2 from (Naik, 2009), Fig 2a from (Laurenti and Göttgens, 2018), Fig 2c from (Naik, 2020)). Importantly for both these models, the precise phenotypic HSPC stage at which these putative patterns of priming or commitment occur is not known.

Here, we attempt the first empirically derived clone-centric landscape of haematopoiesis. By demonstrating that heterogeneous HSPC clonal fate is clone-intrinsic and heritable in vitro, we then assess multiple HSPC stages for their clonal fate in vivo at multiple timepoints, revealing extensive lineage commitment and bias. Last, we integrate these empirically derived stage- and time-series clonal fate data into a novel agent-based clone-centric model of haematopoiesis. This model supports a conceptual framework in which multiple tracks of varying potency are underlying haematopoiesis, and from which clones and their progeny progress and transit as their potency decreases.

## Results

### Evidence of discrete tracks of fate potency at multiple HSPC stages

To better understand the nature of lineage bias amongst Lin^−^Sca1^+^cKit^hi^ (LSK) fractions of cells we developed a new in vitro assay for multipotency and imprinting leveraging our prior clone-splitting approaches (Lin et al., 2018; Naik et al., 2013; Tian et al., 2021). Single HSPCs were first expanded in Stem Cell Factor (SCF) until they reached >25 cells, then daughters were split randomly into multiple parallel tests for fate potential: conditions that differentiate progenitors into B cells, T cells, dendritic cells (DCs) and myeloid cells (Figure 1A, Methods). In this way we gauged clonal multipotency for a total number of 188 clones over two independent experiments. We observed extensive fate heterogeneity (Figure 1B) and, importantly, this was intrinsically programmed as demonstrated by the high conservation of fate between replicates for individual clones (Figure 1C), and when all clones were tested statistically (Figure 1D). Very few HSPCs were multipotent let alone oligopotent, with very few clones able to generate 8 or more cell types out of 12 (Figure 1E). We next repeated this experiment using the index sorting function of the flow sorter, which records the surface phenotype of the individual cells, to link each clone’s potency with a phenotypically defined HSPC population (Jassinskaja et al., 2023). Strikingly, many clones with a similar fate (a cluster within the 1D UMAP) spanned multiple HSPC stages (Figure 1F). One interpretation of this data was that clones with a given fate bias represent discrete populations that are connected developmentally across different HSPC stages e.g. B cell-biased HSCs develop into B cell-biased MPP4s. Therefore, rather than a continuum of ‘low-primed’ states (Hirschi et al., 2017) this was a first hint that HSPC fate biases are more discrete in nature and belong to independent clonal fate trajectories or ‘tracks’ which may traverse classically defined gates but for which potency could already be highly restricted.

**Figure 1.**
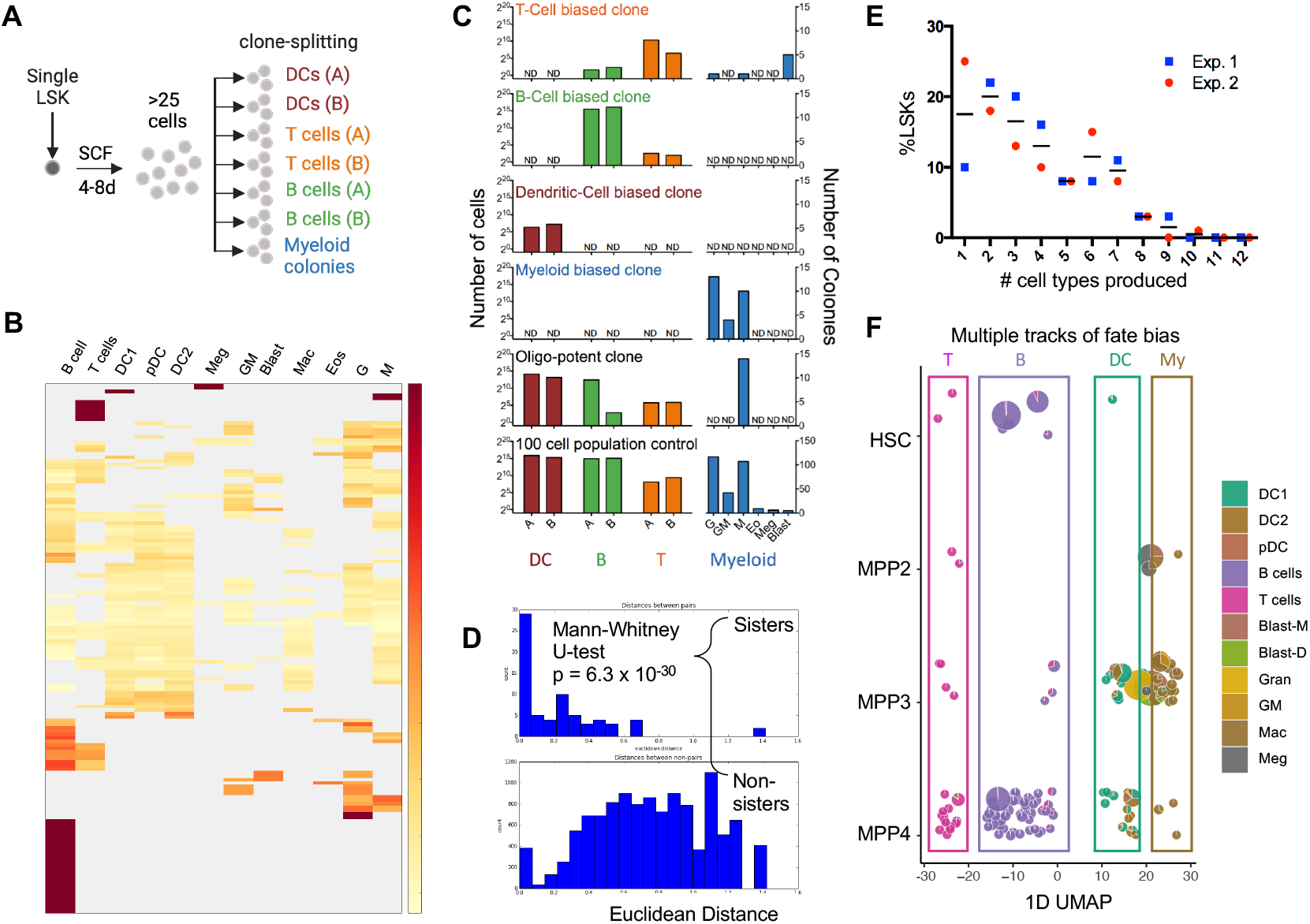
In vitro test of HSPC multipotency infers multiple tracks of lineage bias/commitment. A. Experimental design: single LSKs expanded in SCF with >25 visible progeny were split across 7 conditions to test potency including technical replicates for DC, B cell and T cell fate, and myeloid colony assays. B. Heatmap representation of clonal fate heterogeneity across two experiments. Scale bar indicates proportion of clonal fate towards indicated lineages. C. Fate output of five representative example clones, and a population control, that demonstrate fate heterogeneity between clones but fate concordance within clones. ND, not detected. D. Euclidean distance between sister wells vs random sisters, with indicated p-value. Short distances represent highly concordant fates between sister wells. E. Proportion of LSKs with the indicated potency (# cell types produced). F. LSKs were index sorted by flow cytometry prior to testing as in A. Clones positioned according to population of origin (jittered in y-axis per row for ease of visualisation) and 1D UMAP (x-axis) and represented as pie chart for fate bias. Manual addition of observed ‘tracks’ of indicated lineage bias. LSK, Lin^−^Sca1^+^cKit^+^; SCF, stem cell factor; M, Mac, macrophage colony; GM, granulocyte/macrophage colony; G, Gran, granulocyte colony; Meg, megakaryocytes; Eos, eosinophils; Blast-M, multicentric blast colony; Blast-D, dispersed blast colony.

### A multi-stage barcoding approach allows comprehensive mapping of a haematopoietic landscape

Next we opted to comprehensively characterize the heterogeneity of clonal lineage output of HSPCs in vivo, by developing a multi-stage clonal lineage tracing approach using cellular barcoding (Figure 2A). We purified and barcode-labelled six broad categories of HSPC populations spanning different stages of haematopoiesis, including early multipotent populations (LT-HSC, ST-HSC and MPP) and lineage-restricted progenitors (CLP, CMP and MDP) (Figure S1A). Of note, while the CD150^+^Flt3^−^ gate used to define ‘LT-HSCs’ did harbour CD150^+^ MPP2s, we are unlikely to be measuring any MPP2-derived progeny given the late time points (d56 and d112) and no assessment Mk/E lineages, when compared to known MPP2 fate and kinetics (Pietras et al., 2015). Similarly, the CD150^−^Flt3^−^ ‘ST-HSC’ gate contained MPP3s, but the time points assessed (d28 and d56) excluded much of MPP3-derived progeny, which derive earlier (Pietras et al., 2015).

**Figure 2.**
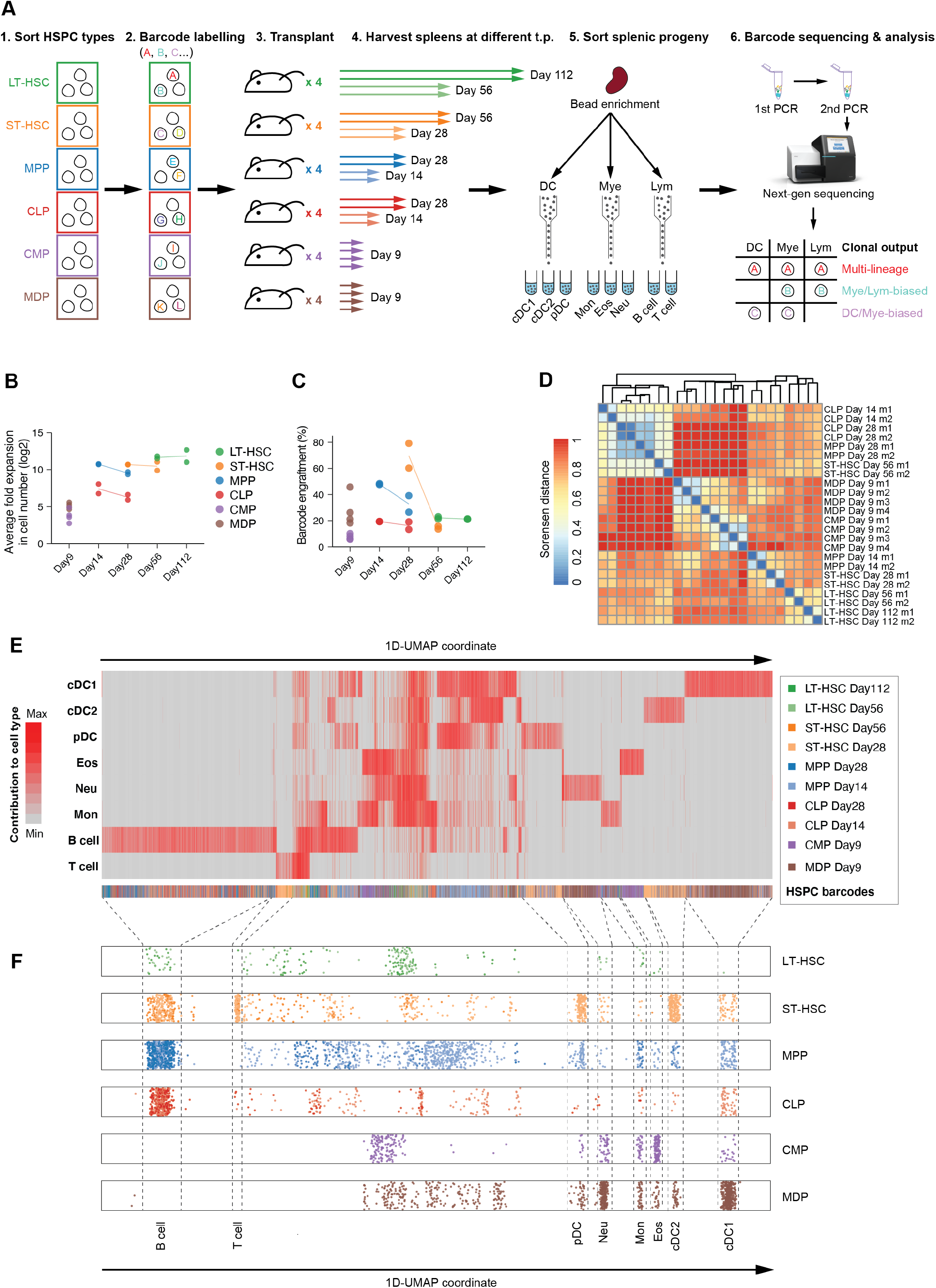
A multi-stage barcoding approach allows comprehensive mapping of clonal lineage commitment of haematopoiesis. A. Experimental set up. Purified HSPC populations were barcode labelled and transplanted into sub-lethally irradiated mice (5 Gy). Spleens were harvested at the indicated time points and progeny cell types were FACS sorted. Barcodes from each cell type were amplified, sequenced and analysed. B. Number of donor-derived cells per recipient over the number of transplanted HSPCs. C. Percentage of barcodes detected per recipient over the number of transplanted barcoded HSPCs. (B, C) Each dot represents one recipient. Colour depicts HSPC group. Line connects the same HSPC group over different time points. D. Heatmap showing Sorensen distance between replicate mice based on barcode lineage fate. Low values indi-cate highly similar patterns of clonal output. E. Heatmap of 5018 barcodes pool from all mice after QC, annotated with HSPC/day groups and ordered by 1D UMAP. Row: contribution to cell types; column: barcode F. Clones ordered by 1D UMAP and separated according to HSPC population of origin. Note multiple tracks of HSPCs with different potencies. LT-HSC, long-term haematopoietic stem cell (HSC); ST-HSC, short-term HSC; MPP, multipotent progenitor; CMP, common myeloid progenitor; CLP, common lymphoid progenitor; MDP, macrophage-DC progenitor; cDC1, conventional dendritic cell type 1; cDC2, cDC type 2; pDC, plasmacytoid DC; Mon, monocytes; Eos, eosinophils; Neu, neutrophils; Mye, myeloid (Mon, Neu, Eos); Lym, lymphoid.

After barcode labelling, between 1-5 x 10^3^ cells of each barcoded HSPC population was transplanted into cohorts of sub-lethally irradiated (5 Gy) mice and multiple cell types from spleen (covering the DC, myeloid and lymphoid lineages) were sorted (Figure 2A, S1B) for barcode sequencing and analysis (Lin et al., 2021, 2018; Naik et al., 2013) at the indicated time points. The realised fate was assumed to largely be imprinted considering the concordance of sibling fate in Figure 1, and prior studies in vitro (Lin et al., 2018) and in vivo (Naik et al., 2013). To fully characterize the developmental dynamics, especially during the early multipotent stages, at least two mice were analysed at two timepoints for LT-HSCs, ST-HSCs, MPPs and CLPs: the first to capture the time of peak production of each HSPC subset (Pietras et al., 2015), and the second to overlap with the peak production of the downstream HSPC stage (Figure 2A).

As expected, we observed higher engraftment and cell expansion in mice receiving early multipotent HSPC populations than late lineage-restricted progenitors (Figure 2B, S1C&D). Nonetheless, robust barcode detection was found in all mice regardless of the population transplanted and timepoints assessed (Figure 2C, S1E). We next compared whether barcoded clones from different recipients exhibited similar or distinct patterns in lineage fate when transplanted with different HSPCs and analysed at different timepoints. We found that mice from the same HSPC/Day group clustered together on the distance heatmap (Figure 2D), indicating biological reproducibility. Consistently, UMAP visualization of barcodes from the same HSPC/Day group showed even mixing from different mice for all groups (Figure S2A). In contrast, barcodes analysed on different days from the same HSPC-transplanted cohort did show a low correlation, except LT-HSC clones (Figure S2B), highlighting the importance of assessing lineage output of the same HSPC populations on different days in our study design.

To gain an overall appreciation of the resulting dataset, we generated a 1D UMAP-ordered heatmap to visualize the quantitative contribution of each individual progenitor (barcodes in columns) to each cell type (rows) (Figure 2E). Consistent with the reported fate heterogeneity in the literature (Lee et al., 2017; Lin et al., 2021; Naik et al., 2013; Notta et al., 2016; Velten et al., 2017; Weinreb et al., 2020), there were notable clusters of fate that were uni-, oligo- and multi-outcome. Importantly, these patterns were often not exclusively derived from a single HSPC subset (annotated on bottom row), indicating the phenotypic definition used to purify the HSPC populations were insufficient to isolate homogenous populations according to their fate profile.

To better visualise whether we could observe similar ‘tracks’ of fate bias that spanned multiple HSPC stages as in the in vitro experiments (Figure 1F), we generated a scatter plot ordering all barcodes according to their clonal fate using 1D-UMAP coordinates on the x-axis, and separating clones by developmental stage (LT-HSC, ST-HSC, MPP, CLP, CMP or MDP) on the y-axis (Figure 2F). Indeed, we found striking patterns of uni-, bi- and reduced potency tracks emerging early in the haematopoietic landscape and remaining consistent throughout the process. Although these data represent snapshots of HSPC fate, they likely represent progenitors that differentiate along separate trajectories or tracks of restricted/biased fate.

### Clonal lineage-biased output is evident from the earliest stages of haematopoiesis

We then characterised the data globally to understand its robustness, and how features of engraftment, potency and clone size differed between HSPC subsets. To examine the extent of fate bias in HSPCs across multiple stages, we first defined each barcode’s clonal potency based on their binary lineage output i.e. how many cell types were generated. For this analysis, the amount of cells produced for each cell type was not taken into account. (Figure 3A). This analysis revealed a strikingly high proportion of lineage-biased clones in all HSPC populations across different timepoints, even within the primitive LT-HSC and ST-HSC compartments. Interestingly, despite ST-HSCs representing an earlier progenitor than MPPs in the classical haematopoietic hierarchy, a higher level of multipotency was observed in MPP clones on day 14 than ST-HSC clones on either day 28 or day 56 (Figure 3A). In fact, no ST-HSC clones on day 28 were found to generate all eight cell types and nearly 80% had restricted fate toward only one cell type.

**Figure 3.**
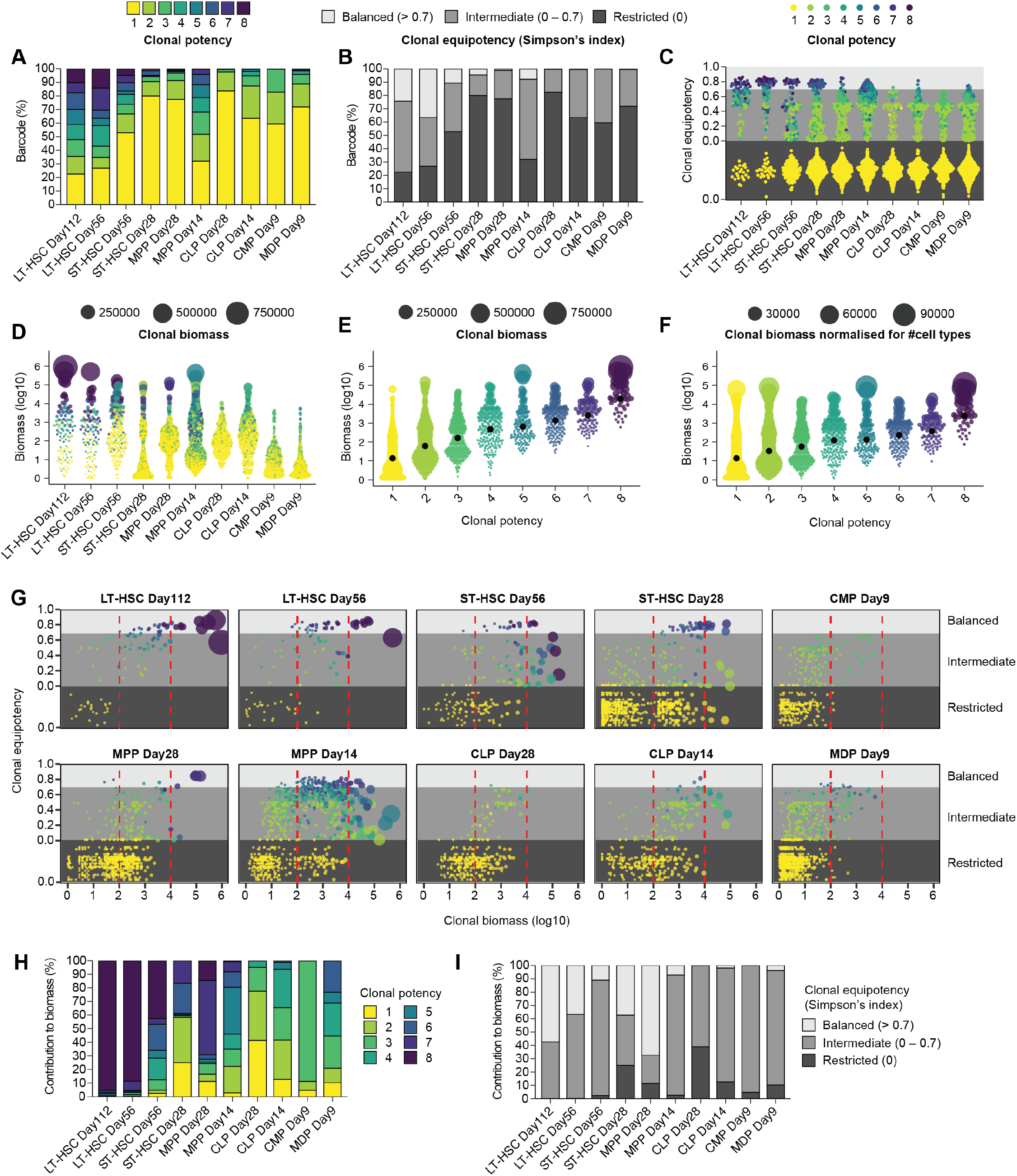
Clonal lineage-biased output is evident from the earliest stages of haematopoiesis. A. Percentage of barcodes with different clonal potency as defined by the number of cell types produced, regardless of cell number generated. B. Percentage of barcodes with different clonal equipotency, as defined by Simpson’s index to measure the degree of balanced lineage output. C. Clonal equipotency of individual barcodes. D. Clonal biomass of individual barcodes from different HSPC/day groups. E. Clonal biomass of individual barcodes with different clonal potency. F. Clonal biomass normalised for number of cell types of individual barcodes with different clonal potency. G. Scatter plots comparing clonal equipotency and clonal biomass from each barcode in each HSPC/Day group. H. Percentage of biomass produced by barcodes with different clonal potency within each HSPC/day group I. Percentage of biomass produced by barcodes with different ranges of clonal equipotency within each HSPC/day group (C-G) Each dot represents one barcode. Colour depicts clonal potency. (D, E & G) Dot size is scaled clonal biomass (linear). (F) Dot size is scaled clonal biomass per cell type (linear).

We next assessed clonal equipotency, which, in contrast to clonal potency, considers how balanced clonal output was across lineages as measured using the Simpson’s index (Figure. 3B&C). For example, if a clone generated equal proportions of every cell type (e.g. 5% of B cells, 5% of monocytes, 5% of neutrophils, etc), it would be considered perfectly balanced (equipotency = 1). If in contrast it only generated a single cell type it would be considered restricted (equipotency = 0). We binned clonal equipotency into three categories including balanced (>0.7), intermediate (>0; <0.7) and restricted (0). We observed strikingly low equipotency across all HSPC/day groups, even in the most primitive LT-HSCs analysed on both day 56 and 112 (average equipotency score < 0.5) (Figure S3A). A positive correlation was found between clonal potency and equipotency, where most clones generating two to five cell types had intermediate equipotency scores between 0 and 0.7, and clones with highly balanced output (equipotency > 0.7) produced at least six cell types (Figure S3B). These balanced clones were rare and were almost exclusively found in the early HSC/MPP groups (Figure 3B&C). In contrast, most clones were found to have a restricted or intermediate output (equipotency < 0.7), even within classically defined LT-HSCs. Together, our results reveal that lineage-biased output is a prominent clonal feature that originates from the earliest stages of haematopoiesis.

### Relationships between clonal potency, equipotency and biomass

We next explored the distribution of clonal biomass (i.e. clone size) and observed differences by orders of magnitudes between the largest and smallest clones within different groups, where the largest clones in the dataset produced close to 1 x 10^6^ cells (Figure 3D). Consistent with prior studies (Lee et al., 2017; Velten et al., 2017), clones with higher potency tended to have larger biomass (Figures 3D&E; S3C). Importantly, this was not due to more cell types being produced, as similar correlation was still found when normalized to the number of cells generated per cell type (Figures 3F & S3D).

To better appraise the interplay between clonal potency, equipotency and biomass, we plotted individual clones according to their clone size (x-axis) and equipotency (y-axis), with visual separation between low (0), intermediate (0-0.7) and high equipotency (>0.7) (grey colour gradient), and between small (<100 cells), medium (100-10^4^ cells), and large (>10^4^ cells) clones (red lines) (Fig. 3G). Dot size was scaled according to clone biomass and coloured according to clonal potency. This visualization revealed an imperfect correlation between any of these three features and that additional complexity and heterogeneity was apparent (Figure 3G). In particular we noted;

- LT-HSC: Although approaching equipotency, no LT-HSC clone had an equipotency of 1, and two of the largest clones were <0.7. Medium sized clones were heterogeneous in their equipotency.
- ST-HSC: While a few ST-HSC clones would be considered truly large and equipotent (top right), the majority were not. Many clones were restricted in their potency, especially at day 28 where many uni- outcome clones were present, including some that were very large.
- CMPs, CLPs, MDPs: Considering these represent restricted downstream progenitors, they had a low potency and were smaller in clone size compared to upstream HSPCs. However, there were many uni- outcome clones in all populations.

Next we quantified the percentage of biomass contributed by clones according to the different degrees of clonal potency given the observation that multi- and equi-potent clones tended to have larger biomass (Figure 3E&F). We found that although most clones were lineage-biased within most HSPC groups (Figure 3A&B) they differed in their relative contribution depending on the HSPC type (Figure 3H&I). The largest contribution from uni-output clones were found in ST-HSCs and CLPs analysed on day 28. In these two groups, more than 80% of barcodes were found to have restricted fate (Figure 3A), which contributed to 20-40% of total biomass for a given lineage (Figure 2H&I). In contrast, although a similar proportion of restricted clones derived from MPPs on day 28, their biomass contribution was much smaller (∼10%) (Figure 3A, H&I). The most striking imbalance was found in the LT-HSC clones, where more than 95% of progeny were produced from only a few LT-HSC clones, and which generated seven or eight cell types (Figure 3A&H). This demonstrates that only a very small fraction of the LT-HSC pool are responsible for the phenotype observed during bulk transplantation experiment of the same population, which has been observed previously (Jordan and Lemischka, 1990; Lu et al., 2011; Naik et al., 2013).

When clones were classified based on equipotency, balanced clones generated approximately half of progenies and the rest were derived from LT-HSCs with intermediate equipotency (Figure 3I). Similarly, balanced and intermediate clones represented the major source of numerical output in most groups. Collectively, our results reveal substantial heterogeneity in clonal features including clonal potency, equipotency and biomass, and highlight the abundance of lineage-biased clones from both the early multipotent and late lineage-restricted stages of haematopoiesis.

### Mapping of clonal lineage output in different stages of haematopoiesis

We next systematically mapped the heterogeneity in clonal lineage output of barcoded HSPCs from different developmental stages. We generated UMAP visualization of all barcodes pooled from all groups, using the number of cells generated per cell type per barcode as input. This resulted in a 2-dimension UMAP plots where barcoded clones (dots) with similar patterns in lineage output were spatially located in proximity (Figure 4A-C). We first annotated these UMAP plots based on clonal contribution to different progeny cell types, which allowed inspection of clonal lineage fate and separation of barcodes into clusters with different fate combination (Figure 4A&B). Unlike uni-output clones that formed separate clusters on the UMAP, a clear distinction was not observed for multi-outcome clones (Figure A&B). This was because barcodes that produced more than one cell type were highly heterogenous in their lineage output, with differential composition and abundance of cell types produced (Figure 4A). Despite this, annotation of regions enriching for certain fate combination was possible (Figure 4B). This was because a clone’s proximity in the UMAP projection to the different uni-output clusters tended to correlate with its potential for the corresponding fate (Figure 4A&B). For example, barcodes located at the bottom half of the UMAP plot all produced B cells, which was in closer proximity to the B-only cluster. In contrast, only a few B cell-producing clones were found to locate at the top half of the UMAP. Amongst these, all were found to locate within a cluster of clones with multi-lineage output (DC ± Mye ± Lym; Figure 4B). Together, UMAP analysis allowed separation of barcoded clones into clusters with different fate combination.

**Figure 4.**
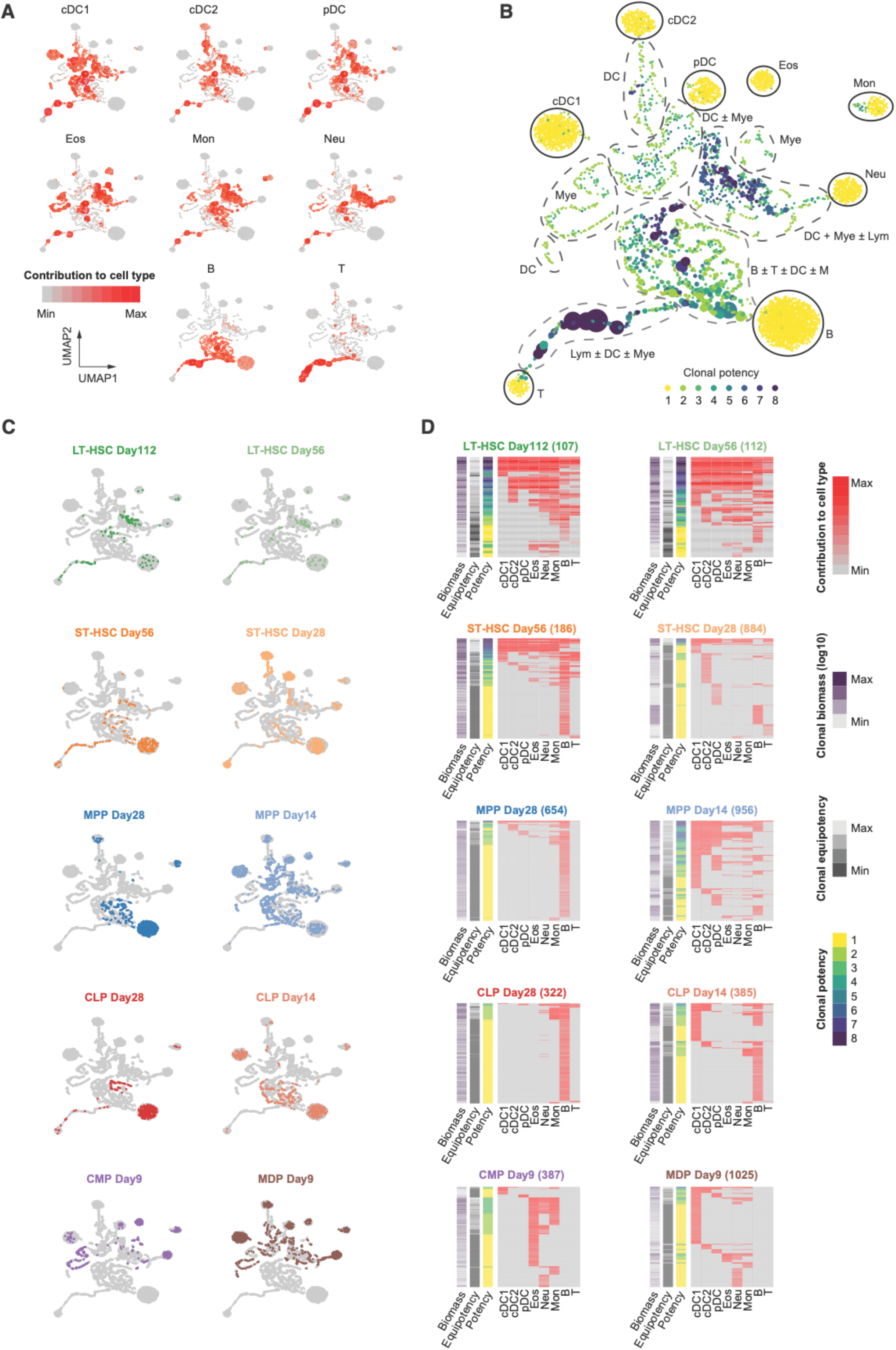
Mapping of clonal lineage output in different stages of haematopoiesis. A. UMAPs colored by contribution to each progeny cell type. Size depicts clonal biomass (linear). B. UMAP colored by clonal potency, with manual annotation of fate clusters. Size depicts clonal biomass (linear). C. UMAPs colored by specified HSPC/day group, with other barcodes colored in grey. D. Heatmaps of barcodes from each HSPC/day group. Each row depicts a barcode. Right columns show contribution to different progeny cell types. Left columns annotate barcode information including clonal potency, clonal equipotency and clonal biomass (log10). Barcodes are ordered by binary fate classification (see Methods).

We next assessed clonal lineage output of different barcoded HSPCs analysed on different time points (Figure 4C). In addition, we generated individual heatmaps showing contribution to cell types (column) by all barcodes (rows) from each HSPC/day group, with annotation of additional barcode information including clonal potency, equipotency and biomass (Figure 4D). Overall, LT-HSC clones exhibited highly similar pattern between early and late time points, with relatively low proportion of uni-output barcodes compared to other HSPC types. Importantly, these were mostly B-only clones. In contrast, clonal behaviour of ST-HSCs was different between day 56 and 28, with only 186 barcodes detected from two mice on day 56, while nearly 5-fold more barcodes were recovered on day 28 (Figure 4D). In addition, most of the uni-output ST-HSC barcodes were B- or T-only on day 56, whereas restricted ST-HSC clones on day 28 had heterogenous output with large numbers of clones producing only DCs, B or T cells (Figure 4C & D). Similarly, MPP output was highly discordant between time points. On day 28, almost all MPP clones were B cell producing, with the majority only generated B cells (Figure 4C & D). This may reflect MPPs containing a population corresponding to the recently discovered lymphoid-primed progenitor (LPP) population (Amann-Zalcenstein et al., 2020; Klein et al., 2022) or clones whose progeny emerge differentially in early (myeloid) vs late (lymphoid) waves of cell production (Naik et al., 2013). Conversely, MPP day 14 barcodes distributed across different regions of the UMAP, indicating the presence of clones with almost all combination of fate detected in the dataset (Figure 4C & D).

### Multiple tracks of haematopoiesis at the clonal level

Motivated by the observation of multiple fates deriving from different HSPC populations varying in potency, balance and output, and across time, we set out to construct a minimal agent-based model of clonal dynamics that can explain the dynamics of fate heterogeneity we observe in our data. The classical “ball and stick” model (Figure 5A) of haematopoiesis (Laurenti and Göttgens, 2018; Naik, 2020) implies a progressive, stepwise restriction of fate over developmental time, where cells within each progenitor compartment or sub-compartments are modelled as homogeneous in their fate potential (see Figure 5C). Under this model, observed variability in fate among clones originating from a common compartment may arise solely from stochastic effects. This model, however, fails to explain the presence of large groups of multipotent as well as fate-restricted (including unipotent) clones observed in our data across different progenitor stages, including LT-HSC and ST-HSC compartments that are conventionally regarded as multipotent.

**Figure 5.**
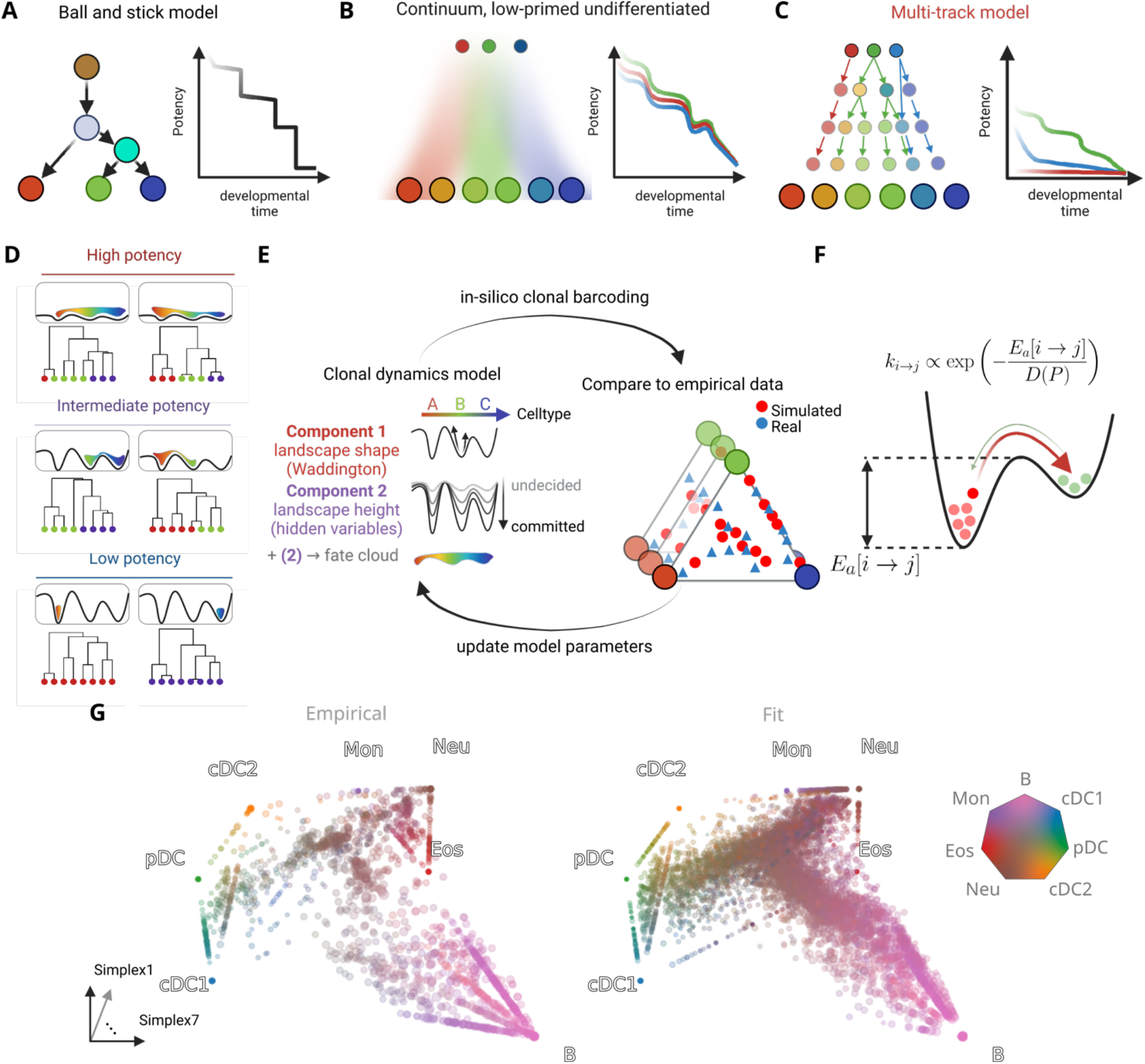
An agent-based mathematical model of clonal haematopoiesis. A. Classical ball and stick model: discrete compartments that do not cannot capture clonal fate heterogeneity within a population and exhibit stage-wise loss of potency. B. Continuous model: continuum of low-primed undifferentiated (CLOUD) HSPCs whose daughters can explore the fate continuum based on exposure to lineage priming regulators. C. Multi-track model: A spectrum of discrete progenitor states whose fate is imprinted and heritable to downstream progenitors, and where lineage commitment can occur at many different stages of haematopoiesis depending on the clone. D. Example clonal “fate clouds” and their corresponding clonal trajectories with examples of high-, medium- and low-potency clones, in a landscape with three coloured cell types. E. Schematic illustrating the agent-based model of clonal fate determination and the iterative procedure for fitting to data. F. In our model, dynamics of clonally related cells are driven by a discrete Waddington’s landscape in which stochastic changes in fate bias are determined by developmental energetic barriers between attractors in the landscape. G. Simplex layouts of clonal profiles observed in the empirical data (left), fitted model output (right)

Here, we propose a mathematical model of clonal dynamics that explicitly accounts for fate heterogeneity among progenitors at each stage of haematopoiesis (Figure 5B). Conceptually, this can be thought of as augmenting a conventional compartmental model (i.e. ball and stick model) of development with a *clone-intrinsic* (i.e. heritable) *variable P* which affects the fate decision making of its constituent cells, and thereby determines clonal potency. The biological relevance of this corresponds to aspects of cellular state that are heritable and are thus shared by all cells deriving from the same progenitor. For instance, these may be features that are not measured or otherwise accounted for in conventional definitions of progenitor cell states (e.g. chromatin modifications, genome topology, methylation) and therefore not directly measured in transcriptomic studies, or alternatively they may be subtle gene expression signatures that cannot be readily uncovered without fate information (MacArthur et al., 2009; Mold et al., 2024; Tian et al., 2021; Weinreb et al., 2020; Zechner et al., 2020).

We construct a minimal model that consists of two main components: (1) a “Waddington’s landscape” which differentiating cells traverse, in which attractors correspond to defined cell fates, and (2) a time-dependent variable *P* which influences clonal potency, controlling the propensity for cells to transition between fate attractors in the landscape. Importantly, *P* is a clone intrinsic property which decreases along developmental time. This can conceptually be understood as an increase in “steepness” of the landscape faced by cells within a given clone as they mature: a flatter landscape (relatively high *P*) at an early stage allows cells to change their state more easily and therefore associated with production of multiple cell types, whereas a steeper landscape (relatively low *P*) at a later stage corresponds to subclones and cells of restricted potency. Note however that in our model high and low values of *P* are possible at any HSPC stage, allowing for both multi- and uni-outcome clones in all stages of development. In Figure 5D, we provide cartoon illustrations of the effects of the landscape steepness on clonal output, for multipotent, oligopotent, and unipotent example clones. In Figure 5D, we provide cartoon illustrations of the effects of the landscape steepness on clonal output, for multipotent, oligopotent, and unipotent example clones.

Our agent-based model implementation simulates the trajectory of proliferating clones, from single progenitor cells at the time of barcoding up to the time its output is measured as mature cell types. Since we only observe the aggregate effects of cell division and death, we capture both by a single, effective, proliferation rate. Note however that cell loss, especially if cell type specific could influence clonal output at different times of sampling; due to the challenge of measuring cell death however this aspect is currently ignored in our model. Simulated clonal output is compared to empirical data using the unbalanced Sinkhorn divergence. To perform model fitting, (i.e. to minimise this divergence), parameters are iteratively updated employing the evolutionary optimisation algorithm CMA-ES (see **Materials and Methods** for details). In Figures 5(E, F) we provide a schematic of our agent-based model and the fitting approach.

To visualise the empirical data as well as the output of our model in the same coordinate system, we used a simplex layout of the space of possible fate outcomes in Figure 5G. In this visualisation, the fate of each clone is represented as a probability vector over the observed cell types. Each vertex in the simplex plot corresponds to a unipotent fate outcome for that cell type. Any possible fate outcome can be therefore expressed as a point lying in the span of the unipotent fates. Any fate outcome can be represented as a list of frequencies of contribution towards each of the 7 celltypes we consider. The space of all possible fate outcomes is therefore 6-dimensional, so we additionally carried out a simple dimensionality reduction using multidimensional scaling (MDS) to obtain a 3-dimensional reduced representation of fate space, which is shown in Figure 5G. Specific patterns of fate bias in the empirical data can be interpreted in this layout. For instance, concentration of points along the “edges” of the simplex between cell types correspond to bipotent clones that were found to produce varying proportions of two cell types, whereas clones found to lie in the interior of the simplex correspond to oligo- or multipotent clones. We find that the simulated distribution of fates from the fitted model recapitulates key aspects of the empirical data. We observe for instance a trifurcate structure between myeloid, DC, and B-cell lineages, as well as the presence of bipotent clones among the DC and myeloid lineages - notably the close relationship of monocytes with neutrophils, and cDC1s to cDC2s.

In Figure 6 (A, B), we compare the simulated model output to the empirical data using a unidimensional projection of the fate space simplex against developmental time. This illustrates the successive restriction of fate in the model towards unilineage tracks (solid colours). At the earliest point in developmental time there is a concentration of multipotent progenitors, reflected as a density of brown points lying outside unipotent “tracks”. We note the presence of a well-delineated B-cell track (shown in pink) that reflects the presence of unipotent B-lineage fates from the LT-HSC progenitor pool in the empirical data that grows over developmental time. As developmental time progresses, the gradual emergence and growth of unipotent tracks can be observed, such that eventually all fate outcomes are confined to one of the unilineage tracks. In Figure 6B, we overlay the empirical data from each labelled progenitor stage over the model outputs, demonstrating that current phenotypic definitions of HSPC populations are constrained by conceptual limitations and highlighting the need for better measures and prospective markers of lineage potency.

**Figure 6.**
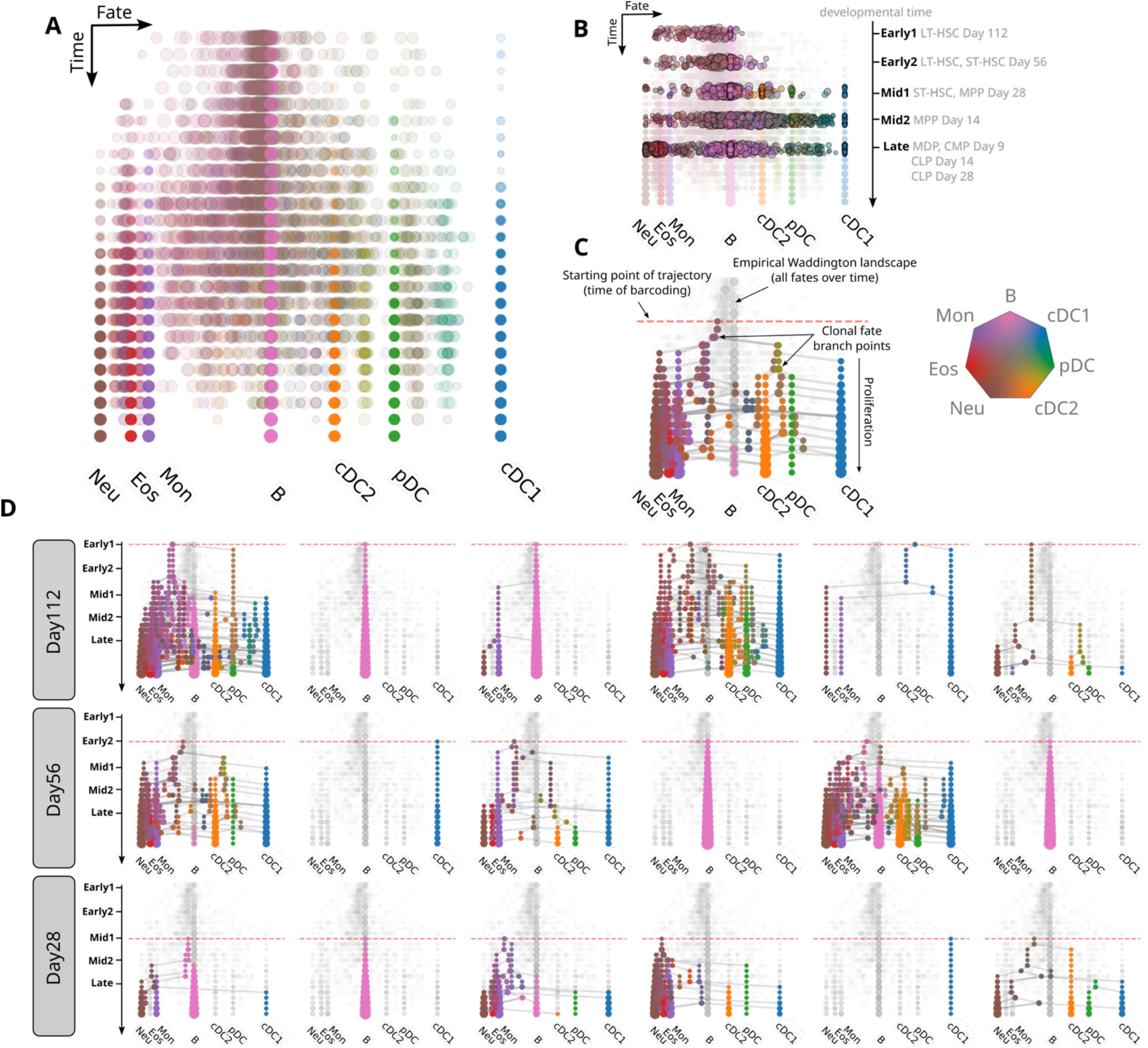
A clone-centric landscape of haematopoiesis. A. Empirically derived landscape of haematopoiesis represented as simulated fates from clone-centric agent-based model, projected onto a unidimensional layout derived from the simplex plot of Figure 5 against developmental time. B. Empirical barcoding data overlaid on the coordinates of the empirically derived landscape in (A), split up across developmental time by progenitors and experimental timepoint. C. Annotated example of a simulated clonal trajectory shown as a tree. Dots represent sub-clonal populations of cells at a given developmental time (y-coordinate) and with a certain fate potential (x-coordinate). Dot size corresponds to log-cell number. Lines correspond to changes in fate potential over time. D. Sampled clonal trajectories from our fitted agent-based model, corresponding to multipotent, oligopotent and unipotent fate outcomes at different stages of clonal trajectory initiation.

Finally, we simulated trajectories of single clones from our model in the form of trees, starting from different developmental stages. We show an annotated example of such a clonal lineage tree in Figure 6C in the 1-dimensional simplex coordinates versus time. In Figure 6D, we show more sampled trees starting from different points in developmental time. We observe among the simulated clonal trajectories that there is significant variability in both the final readout, as well as the intermediate, developmental “kinetics” of the trajectory. In particular, we observe that a fraction of trajectories, even from day112 (corresponding to the LT-HSC compartment), are confined to the B lineage.

## Discussion

Our revised multi-track landscape of haematopoiesis is a departure from graphical estimations of the process of haematopoietic differentiation, and single cell ‘omics inferred ‘trees’ or pseudotime trajectories. Instead, our model provides an empirically derived representation of when and where in the haematopoietic landscape a single HSPC commences its trajectory according to its fate potential, where its daughters may branch into new tracks of reduced potency, and the precise cellular output in terms of cell numbers of intermediate progenitor and progeny cell types. Importantly, the model a) allows the allocation of historically defined HSPCs to this landscape, b) is built on the systematic assessment of clonal multi-lineage fate in vivo rather than in vitro, and c) implements novel agent-based modelling approaches to provide a clone-centric rather than cell-centric perspectives.

We demonstrate that commitment of single HSPCs to tracks of reduced potency can occur at multiple points along the haematopoietic landscape, a significant deviation from the ball-and-stick model. In many cases this commitment occurs much earlier than might have been anticipated, certainly when considering the classic models of haematopoiesis. Even the more recent ‘low-primed undifferentiated HSPC’ continuous models where the implicit assumption that the majority of cells retain a certain degree of plasticity in their lineage contribution (Hirschi et al., 2017; Laurenti and Göttgens, 2018; Velten et al., 2017), is not supported by our data. Instead, we observe evidence of extensive hardwired and discrete lineage bias at a clonal level amongst HSPCs. Therefore, we propose ‘multi-track’ as a more intuitive and informative term that distinguishes the concepts of a flexible continuum of states (Hirschi et al., 2017; Velten et al., 2017), from a (potentially numerous) set of pre-determined paths that exist in parallel within HSPC populations (Graphical abstract, Figure 5c, adapted from Fig 2 from (Naik, 2009), Fig 2a from (Laurenti and Göttgens, 2018), Fig 2c from (Naik, 2020)).

Whether suitable surface markers or reporters will be discovered in the future to prospectively isolate such cells that belong to a certain track e.g. lymphoid-committed (ST-HSCs to MPPs to CLPs), remains to be seen. Certainly, our original description of fate heterogeneity amongst MPP4s into lymphoid-, DC- and myeloid-biased LMPPs (Naik et al., 2013) was not orthogonally validated until prospective isolation of lineage biased progenitors using reporters for *Dach1* (Amann-Zalcenstein et al., 2020) and *Dntt* (Klein et al., 2022) for lymphoid commitment, and *Irf8* (Kurotaki et al., 2019) for DC commitment. One may predict that factors that define fate heterogeneity in the present study will also take time to be discovered by testing fate of HSPC subpopulations separable by TFs and epigenetic mechanisms (Buenrostro et al., 2017; Giladi et al., 2018), immunophenotypic definitions (Triana et al., 2021; Zhang et al., 2024), or factors that are identified by relating the transcriptional state of parental cells to their destined fate using SIS-seq (Luyi et al., 2018; Tian et al., 2021) or similar approaches (Biddy et al., 2018; Emert et al., 2021; Rodriguez-Fraticelli et al., 2020; Weinreb et al., 2020). Identifying such discrete populations in different tracks does not exclude the importance and role of microenvironment in instructive versus permissive establishment of fate.

It is also important to note that this study employed isolation and transplantation into myeloablated hosts and this is unlikely to reflect steady-state haematopoiesis. Native cellular barcoding technologies are only emerging and population-specific induction of barcode generation does not yet extend beyond Cre-induced models (Feng et al., 2024; Pei et al., 2020; Weber et al., 2023). Even with such models, identifying an appropriate driver for the literature-defined stem and progenitor populations remains contentious. To that end, prospective isolation, barcoding and transplantation of well-characterized populations is currently the only means to determine clonal fate profiles, followed by integration of cell fate data, that we achieved in the present study using agent-based modelling.

Longitudinal studies are also an experimental ideal. However, while we and others have examined the clonal dynamics of HSPC fate to a restricted set of cell types in vitro (Karamitros et al., 2018; Lee et al., 2017; Lin et al., 2018; Notta et al., 2016; Velten et al., 2017), measuring the potency for all possible lineages can only be achieved in vivo, where longitudinal sampling of bone marrow (the major site of lineage commitment) is not feasible.

Mathematical models of hematopoiesis have a long history (Busch et al., 2015; Kucinski et al., 2024; MacLean et al., 2013; Manesso et al., 2013; Olariu and Peterson, 2019; Peixoto et al., 2011; Pellin et al., 2019a; Perié et al., 2014), yet to our knowledge no existing framework can account for the striking heterogeneity in potency we observe for clones at all HSPC stages. To fill this gap and enhance interpretability of our data we formulated a novel agent-based clone-centric model of blood development at the single cell level, able to integrate multi-stage and multi-time point fate data into an empirically derived discrete Waddington’s landscape of haematopoiesis. For genuine longitudinal clonal trajectories, however, future studies could either optimise in vitro models to permit multi-lineage haematopoiesis (Lee et al., 2017; Notta et al., 2016; Weinreb et al., 2020) thereby allowing serial sampling of clonal trajectories across real time (versus single cell RNA-seq inferred pseudotime) or through the recording of lineage commitment events through cell history recorders (Masuyama et al., 2022).

How these trajectories are skewed in conditions of demand-adapted haematopoiesis and oncogenic transformation will also be important to consider. For example, can a given oncogene or inflammatory event transform all trajectories equally, or is there be a pre-requisite trajectory that is susceptible to a given transformation (Lin et al., 2021; Tian et al., 2021). This could be addressed using SIS-skew, which links a clone’s normal fate with its perturbed fate (Tian et al., 2021). Lastly, testing these models in human haematopoiesis (homeostatic, during ageing, and in instances of clonal haematopoiesis) will be challenging but necessary; either utilising in vitro or humanised models of human haematopoiesis (Lee et al., 2017; Notta et al., 2016), humanized mice, or inference using natural barcoding: (Lee-Six et al., 2018; Ludwig et al., 2019; Miller et al., 2022; Weng et al., 2024).

In sum, viewing haematopoiesis through a clonal lens, and in the context of a multi-track landscape, reconciles conflicting models and data of lineage-commitment and priming at different stages of haematopoiesis. Future studies could explore whether such a model holds true for the development of other tissues, or apply similar approaches to understanding cancer development.

## Methods

### Mice

All mice were bred and maintained under specific pathogen-free conditions, and protocols were approved by the WEHI animal ethics committee (AEC2018.015, AEC2018.006, AEC2014.031). Mice aged between 8-16 weeks were used. CD45.1 (C57BL/6 Pep^3b^) mice were used as donors and CD45.2 (C57BL/6) mice were used as recipients in the transplantation experiment.

### Flow cytometry

Cell sorting was performed on a BD FACSAria-II/III (BD Biosciences). Data analysis was performed using FlowJo 9.9.6 (Treestar).

### Isolation of BM HSPC populations

Bone marrow cells from ilium, tibia and femur were collected by flushing with FACS buffer (PBS containing 0.5% FBS and 2 mM EDTA) through a 22-gauge needle. Cells were then stained with cKit-APC antibody (In house, 2B8, 1:800) in FACS buffer for 30 minutes at 4 °C, washed and incubated with anti-APC magnetic beads (Miltenyl Biotec cat# 130-090-855; 200 µL beads per 5 x 10^8^ cells per ml) in FACS buffer for 15 minutes at 4 °C. Magnetic-Activated Cell Sorting (MACS) enrichment was performed following the manufacturer’s protocol (Miltenyl Biotec) using LS columns. The cKit-enriched fraction was then stained with a cocktail of antibodies in FACS buffer for 30 minutes at 4 °C including Sca1-A594 (In house, E13-161.7, 1:300), CD150-BV421 (Biolegend, TC15-12F12.2, 1:200), CD16/32-FITC (in-house, 2.4G2, 1:2000), CD11b-BB700 (BD, M1/70, 1:600), IL7Ra-biotin (In house, A7R34, 1:200) and Flt3-PE (eBioscience, A2F10, 1:100). Next, cells were stained with streptavidin-PE/Cy7 (BioLegend, 1:500) in FACS buffer for 30 minutes at 4 °C. Lastly, cells were resuspended in FACS buffer containing Propidium iodide (PI, 1:1000) to exclude dead cells prior to cell sorting.

After exclusion of PI^+^ non-viable cells and doublets based on FCS and SSC, BM HSPC populations were defined as the following: LT-HSCs (CD11b^−^IL7Rα^−^cKit^+^Sca1^+^Flt3^−^CD150^+^), ST-HSCs (CD11b^−^IL7Rα^−^cKit^+^Sca1^+^Flt3^−^ CD150^−^), MPPs (CD11b^−^IL7Rα^−^cKit^+^Sca1^+^Flt3^+^CD150^−^), CLPs (IL7Rα^+^Flt3^+^), CMPs (CD11b^−^IL7Rα^−^cKit^+^Sca1^−^Flt3^−^CD16/32^int^), MDPs (cKit^+^Sca1^−^Flt3^+^ CD16/32^int^). The gating strategy of BM HSPC populations is shown in Figure S1a and we acknowledge these are not pure populations in some cases (see main text).

### Potency assay

We developed a multipotency assay involving the deposition and expansion of clones, following by splitting of daughters into multiple parallel conditions for fate assessment. Viable cKit-enriched were stained for cKit, Sca1, CD150, CD48, Flt3, CD11b and IL7Rα, gated for cKit^+^Sca1^+^CD11b^−^IL7Rα^−^ cells and then single cell index sorted into 96-well round-bottom plates containing StemSpan™ SFEM II (Stemcell Technologies) supplemented with 50 ng/mL stem cell factor (in-house) and 1:10,000 Flt3 ligand (BioXcell), penicillin and streptomycin. During 4-8 days of culture clones were scored daily and those with >25 cells were split the following day by gently mixing the cells with the supernatant. Equal proportions were split in duplicate over 4 different culture conditions. 1. DC culture (200 μl DC conditioned media (supernatant of day 3 Flt3L BM cultures in RPMI 1640 medium) with 1/1600 freshly added Flt3L per well). 2. B Cell cultures (200 µL RPMI 1640 with 1/10000 Flt3L, 1/2000 IL-7 (in-house) and 2000 OP9 cells per well). 3. T Cell culture (200 µL RPMI 1640 with 1/10000 Flt3L, 1/2000 IL-7 and 2000 OP9-DL1 cells per well). 4. Myeloid colony assay (1 mL semisolid agar cultures in Dulbecco’s modified Eagles medium (DMEM) containing 20% newborn calf serum (FCS) in stem cell factor (SCF) 100 ng/ml, erythropoietin (EPO) 2 U/ml, and interleukin-3 (IL-3) 10ng/mL)

#### Cell staining of different subtypes

Dendritic, and B-cell cultures were analysed on day 18 or 19, 28 or 29 post single cell sort, respectively. Fluorescently labelled counting beads (BD Biosciences) (PE for technical replicate A (TR-A) or APC for TR-B) were added prior to staining and cells were subsequently stained with the following panels. DC’s: CD11c-APC (in-house, N418, 1:800), CD172a-A594 (SIRPα, in-house, P84, 1:425), MHC-II-APC/Cy7 (BioLegend, M5/114.15.2, 1:3000) Siglec-H-PE (eBiosciences, eBio440c, 1:1000), CD24-BV711 (BioLegend, M1/69, 1:2500) and CCR9-PE/Cy7 (eBioscience, eBioCW1.2, 1:1000) as well as CD45.2 in either BV421 (BD, 104, 1:3000) (TR-A) or Alexafluor 700 (In house, S450, 1:400) (TR-B). cDC1 were scored as CD11c^+^MHC-II^+^CD24^+^CD172a^−^, cDC2 as CD11c^+^MHC-II^+^CD24^−^CD172a^+^, and pDC as CD11c^+^MHC-II^+^SiglecH^+^CCR9^+^. B-cells: CD19-APC (in-house, ID3, 1:400), B220-APC/Cy7 (BioLegend, RA3-6B2, 1:800), CD4-PE (in-house, GK1.5, 1:400), CD8α-A594 (in-house, 53-6.7, 1:400) and CD3ε-PE/Cy7 (BioLegend, 145-2C11, 1:1000) as well as CD45.2 in either BV421 (TR-A) or Alexafluor 700 (TR-B) and scored as CD19^+^B220^+^ cells. After washing technical replicates were pooled. T-cell cultures were analysed on day 41 or 42 post single cell sort and were stained with: CD19-PE (in-house, ID3, 1:400), B220-APC/Cy7 (BioLegend, RA3-6B2, 1:800) CD4-A594 (in-house, GK1.5, 1:400), CD8α-A647 (in-house, 53-6.7, 1:400) and CD3ε-PE/Cy7 (BioLegend, 145-2C11, 1:1000) and scored as CD3ε ^+^ cells. All cells were analysed on BD LSR Fortessa. For myeloid formation, colonies were fixed, dried onto glass slides, and stained for acetylcholinesterase, Luxol fast blue and hematoxylin, and the number and type of colonies were determined by microscopy (Nikon Eclipse E600, 10x objective).

### Barcode transduction

Barcode transduction was performed as described previously (Naik et al., 2013). Freshly isolated HSPC populations (LT-HSC, ST-HSC, MPP, CLP, CMP, MDP) were resuspended in StemSpan medium (Stem Cell Technologies) supplemented with 50 ng/mL stem cell factor (SCF; generated in-house by Dr Jian-Guo Zhang) and transferred to individual wells of a 96-well round bottom plate at less than 1 × 10^5^ cells/well. Small amount of lentivirus containing the barcode library (8 μL per well; pre-determined to give 5-10% transduction efficiency) was added and the plate was centrifuged at 900 *g* for 90 minutes at 22 °C prior to incubation at 37 °C and 5% CO_2_ for 4.5 hours. After incubation, cells from individual wells were transferred to 10 mL tubes and washed using 8 mL of FBS-containing buffer (10% FBS in PBS or RPMI) twice to remove residual virus. Cells were then washed once using PBS to remove FBS, prior to resuspension in PBS for transplantation.

### Transplantation

Barcode-transduced HSPC populations from CD45.1 BM were transplanted into sub-lethally irradiated (7 Gy) CD45.2 recipient mice. 1,323 LT-HSCs, 4,286 ST-HSCs, 4,762 MPPs, 5,002 CLPs or 4,999 CMPs were injected into four recipient mice per HSPC population, while two recipients received 5,001 MDPs intravenously. In addition, 5,000 CMPs or 5,000 MDPs were transplanted into two recipient mice per population via intra-splenic injection. Spleens of recipient mice were harvested at the indicated time points (Figure 1A).

### Isolation of splenic populations

Spleens were mashed with FACS buffer through 70 μm cell strainers with 3 mL syringe plungers. Red blood cells were lysed by incubating with Red Cell Removal Buffer (RCRB; NH4Cl; generated in-house) for 1– 2 minutes, followed by washing and resuspension with FACS buffer. Cells were first stained with a mixture of antibodies including CD45.1-BV650 (BD, A20, 1:200), CD45.2-PE (BioLegend, 104, 1:800), CD11c-APC (BD, HL3, 1:400) and CD11b-biotin (In house, M1/70, 1:400) at 4 °C for 30 minutes, followed by incubation of anti-APC beads (Miltenyi Biotec, cat# 130-090-855) at 4 °C for 30 minutes. MACS enrichment was performed according to the manufacturer’s protocol using MS columns. The CD11c-APC-enriched fraction (DC fraction) was then stained with F4/80-A700 (In house, F4/80, 1:400), Bst2-Pacific blue (In house, 120G8, 1:200), SIRPα-PEDazzle594 (BioLegend, P84, 1:1000) and CD8α-PE/Cy7 (BioLegend, 53-6.7, 1/400) at 4 °C for 30 minutes. The CD11c-APC-depleted fraction was further incubated with anti-biotin beads (Milteniy Biotec, cat# 130-090-485) and streptavidin-PE/Cy7 (BioLegend, 1:500), followed by another MACS enrichment using MS columns. The CD11b-biotin-enriched fraction (Myeloid fraction) was stained with F4/80-A700 and Gr1-Pacific blue (In house, RB6-8c5, 1:400); while the CD11b-biotin-depleted fraction (lymphoid fraction) was stained with CD19-A700 (In house, 1D3, 1:200) and CD3-Pacific blue (In house, 17A2, 1:400). After the final stain, cells in DC, myeloid and lymphoid fractions were resuspended in FACS buffer containing PI (1:1000) prior to cell sorting.

After exclusion of PI^+^ non-viable cells and doublets based on FCS and SSC, splenic progeny populations were defined as the following: cDC1s (F4/80^low/–^Bst2^−^CD11c^+^CD8α^+^SIRPα^−^), cDC2s (F4/80^low/–^Bst2^−^CD11c^+^CD8α^−^ SIRPα^+^) and pDCs (F4/80^low/–^Bst2^+^ CD11c^int^) from the DC-enriched fraction; eosinophils (eos, F4/80^−^ CD11b^+^Gr-1^int^SSC-A^hi^), monocytes (mon, F4/80^−^CD11b^+^Gr-1^int^SSC-A^low^) and neutrophils (neu, F4/80^−^ CD11b^+^Gr-1^hi^SSC-A^int^) from the myeloid-enriched fraction; B cells (CD11c^−^CD11b^−^CD3^−^CD19^+^) and T cells (CD11c^−^CD11b^−^CD3^+^CD19^−^), CD45.1^+^CD45.2^−^ donor-derived cells within each population were sorted for barcode amplification and sequencing. The gating strategy of splenic populations is shown in Figure. S1b.

### Barcode amplification and sequencing

PCR and sequencing were performed as described previously (Naik et al., 2013). Briefly, sorted splenic populations were lysed in 40 μL lysis buffer (Viagen) containing 0.5 mg/ml Proteinase K (Invitrogen) and split into technical replicates. Barcodes in cell lysate were then amplified following two rounds of PCRs. The first PCR amplified barcode DNA using common primers including the TopLiB (5’ – TGC TGC CGT CAA CTA GAA CA – 3’) and BotLiB (5’ – GAT CTC GAATCA GGC GCT TA – 3’). The second PCR introduced an 82-bp well-specific 5’ end forward index primer (384 in total) and an 86-bp plate-specific 3’ reverse index primer (8 in total) to each sample for later de-multiplexing *in silico*. The sequences of these index primers are available upon request. Products from second round PCR with index primers were run on a 2% agarose gel to confirm a PCR product was generated, prior to being cleaned with size selected beads (NucleoMag NGS) according to the manufacturer’s protocol. The cleaned PCR products were pooled, and sequencing was performed on the Illumina MiSeq platform.

### Barcode data processing and filtering

First, number of reads per barcode from individual samples was mapped to the reference barcode library, which contains 2608 actual unique DNA barcode sequences (available upon request) and counted using the *processAmplicons* function from edgeR package (Version 3.28.0) (Dai et al., 2014; Robinson et al., 2009). Next, low-quality samples were removed, including those with total barcode read counts of less than 500 and those with very low Pearson correlation between technical replicates (0.2). To remove low quality barcodes in the remaining samples, read counts were set to zero for barcodes with reads in one but not the other technical replicate. After this, read counts of each barcode from technical replicates were averaged and normalized to the number of cells, estimated based on cell counting. The clonal biomass (sum of number of cells from all splenic population per barcode) was calculated and barcodes with biomass less than one cell was removed. Lastly, to ensure reproducibility of biological findings, a UMAP density-based filtering step was developed. Briefly, individual UMAP was generated for each group of barcodes (sampled from recipients transplanted with the same HSPC type and harvested on the same day). Barcodes in areas with no overlap between biological replicates were removed.

### Barcode data analysis and visualization

Clonal potency was defined by calculating binary fate of each barcode with regard to the number of cell type produced, using a threshold of 0.01% for each cell type. For example, if the number of cDC1 produced by a barcode was 0.02% of its total clonal biomass/size, this barcode is defined as cDC1 producing. If all eight cell types assessed were more than 0.01%, the clonal potency of this barcode would be 8. Clonal equipotency was defined by computing Simpson diversity of each barcode using the Vegan package (Version 2.6-4). Heatmaps were generated using the pheapmap package (Version 1.0.12). 2d- and 1d-UMAP coordinate for each barcode were computed using the umap package (Version 0.2.10.0). Scatter plots and violin plots were generated using ggplot 2 package (Version 3.4.4).

### Computational modelling

Clonal contributions towards celltypes of interest were quantified in terms of biomass as per (**Barcode data processing and filtering**). Following this, barcodes with a total biomass of less than 5 units were removed prior to downstream analysis and modelling. As described in the main text (see **Multiple tracks of haematopoiesis at the clonal level**), we disregarded contributions to the T lineage due to poor contribution by lentiviral transduced MPPs (Naik et al., 2013), and so contributions to the cDC1, cDC2, pDC, eosinophil, monocyte, neutrophil and B-cell lineages were retained. Barcodes captured from each of the progenitor subpopulations and timepoints are further grouped into five stages prior to modelling, which for convenience are referred to as **Early1**, **Early2**, **Mid1**, **Mid2**, and **Late**.

| Stage | Subpopulations |
| --- | --- |
| **Early1** | LT-HSC (Day112) |
| **Early2** | LT-HSC (Day56), ST-HSC (Day56) |
| **Mid1** | ST-HSC (Day28), MPP (Day28) |
| **Mid2** | MPP (Day14) |
| **Late** | ProDC (Day9), CMP (Day9), CLP (Day14, 28) |

Due to the exponential nature of cell numbers in proliferating populations, we find that absolute biomass often varies by several orders of magnitude across barcodes and cell types. Barcode biomasses are therefore transformed into an estimated division number by applying a log(*x* + 1)-transform, followed by normalising each barcode’s logarithmised cell-type contributions to sum to unity. The output of this pre-processing scheme represents each barcode is represented by a vector of length 7 summing to 1, i.e. a point in the (7-1)-dimensional probability simplex, where the *i*th entry of the *j*th vector corresponds to the normalised contribution of clone *j* to lineage *i*, in the log-domain.

#### Theory

To model the clonal dynamics observed in our barcoding data, we designed a stochastic generative model that describes the evolution of clones over developmental time. To do so, clones are modelled as being a group of cells arising from a common progenitor that evolve over a discrete landscape that delineates cell type. Inspired by Waddington’s analogy of an epigenetic landscape (Waddington, 2014), we choose to model cell dynamics using a dynamical system driven by a potential function. We refer the interested reader to (Zhou et al., 2012) and (Brackston et al., 2018) for a mathematical treatment of potential landscape models of developmental biology.

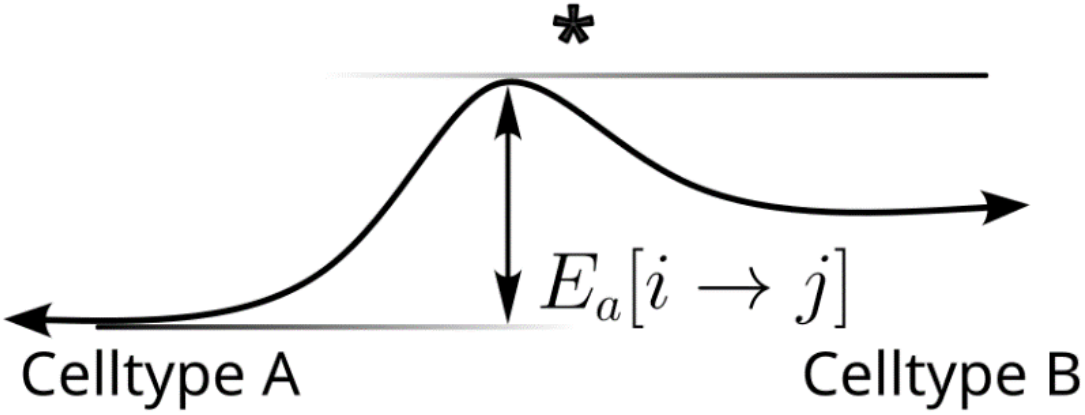

#### Schematic diagram of metastable landscape model

We consider a landscape in which transition paths between any pair of states (for instance, A → B) must pass through an intermediate transition state, denoted here by an asterisk (*). The activation energy barrier, denoted Ea, determines the likelihood of the transition.

Abstractly, define 𝒳 to be a cell state space, and let Ψ(*x*) be a potential function with *k* attractors with each attractor corresponding to a mature cell type. Inspired by the transition state theory of molecular reactions, we assume that each pair of basins (*i*, *j*), 1 ≤ *i* ≠ *j* ≤ |𝒳| is separated by an energetic barrier in the potential landscape Ψ. We will refer to the saddle node *x*^*^ as the *transition state*.

We reduce this continuous model of cell dynamics to a discrete model, in which we identify attractors in the epigenetic landscape with discrete cell types. Let 𝒳 = {*x*_cDC1_, *x*_cDC2_, *x*_pDC_, *x*_Eos_, *x*_Mon_, *x*_Neu_, *x*_B_} be the discrete cell state space. Based on this reduction of the state space, we also reduce the potential landscape Ψ, in which transitions between any two attractors is determined by an energetic barrier. This enables us to obtain a discrete, stochastic dynamical model that can be fit to experimental data.

Let Ψ*_i_* be the potential energy of cell state *x_i_*, then for any other state *x_j_*, a cell-state transition *x_i_* ↔ *x_j_* must proceed through an intermediate state 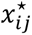 with potential energy 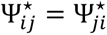. We emphasise that the landscape assumption imposes that the forward and reverse transitions must proceed through the *same* intermediate state. Kramer’s law (Berglund, 2013) provides a rigorous way to derive transition rates between attractors in the setting of a continuous landscape – for our purposes, it will serve to inspire the functional form of the transition rate function. The rate of any transition *x_i_* → *x_j_* is determined by its associated *activation energy* E*_a_*[*x_i_* → *x_j_*] that must be overcome. In our context, this is given by

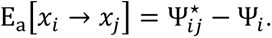

As long as the Ψ*_i_*, 1 ≤ *i* ≤ |𝒳| are not all constant (i.e. all minima are not at the same potential value), this model allows for *asymmetric* transition rates between attractors. The transition rate is then given by

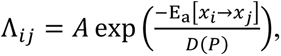

where the diffusivity *D* is a function of the clone-intrinsic potency variable *P* (see **Multiple tracks of haematopoiesis at the clonal level**).

For given rates 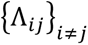 a continuous time Markov chain (CTMC) on the set of attractor states can be constructed: let Λ be the matrix of pairwise transition rates, with Λ*_ii_* = 0 and ***D*** = diag(**Λ1**), where we have written **1** denoting the |𝒳| × 1 vector of all ones. Then the CTMC has generator ***L*** = ***D*** − **Λ**. In a time interval *τ* > 0 therefore, the corresponding transition matrix is calculated as a matrix exponential ***T***_***τ***_ = exp(−*τ**L***).

In addition to the landscape dynamics, proliferation is essential to model clonal dynamics. We use a simple model of cell proliferation – to each attractor *x_i_* we assign a branching rate *β_i_*, and the corresponding division time is exponentially distributed with mean *β*_*i*_^−1^. For such a process with branching rate *β* with *N*_0_ individuals at time *t* = 0, at time *t* = *τ* the number of offspring is a random variable with a negative binomial distribution (Ross, 2010).

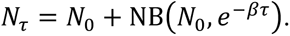

#### Computational implementation

We choose to simulate on a temporal domain of 0 ≤ *t* ≤ 1. On this domain, clones from Early1, Early2, Mid1, Mid2 and Late were simulated starting from *t* = 0, 1/6, 2/6, 3/6 and 4/6, respectively, up to *t* = 1. In practice we discretised the temporal domain into 30 timesteps, i.e. we simulate in timesteps of *τ* = 1/30. Each clone was initialised at its starting time with a single cell in a randomly chosen attractor, chosen according to an initial distribution ***p***_***0***_. We simulated the trajectory of a clone by iteratively applying the temperature-dependent transition kernel, followed by sampling numbers of offspring using the negative binomial model.

To calculate transition rates, we set the prefactor *A* = 1 and set *D*(*P*) = *P*. Furthermore, we chose *P*(*t*; *P*_0_) = (1 − *t*)*P*_0_ + *t P_min_*, where *P_min_* = 0.05 is a small, positive minimum temperature chosen to avoid numerical instabilities. The initial value *P*_0_for each clone is sampled from a distribution *q*_***T***_. We choose an exponential temperature distribution, ***q**_T_* = *P_min_* + Exp(1).

#### Algorithm 1 (Forward simulation)

Inputs: time step *τ*, clone-intrinsic potency schedule *P*(*t*; *P*_0_), prefactor *A*

- Sample initial clone state, ***x***_***0***_ = **1***_X_*_0_ where ***p***_0_ ∼ ***p***_***0***_ and initial temperature *P*_0_ ∼ *q*_***T***_
- *t* ← 0
- **while** *t* ≤ 1:

- Get current temperature *P*(*t* ; *P*_0_)
- Calculate rate matrix **Λ** where 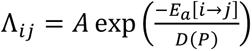
- Calculate transition matrix ***T***_***t***_ = exp(−*τL*) where *L* = diag(Λ**1**) − Λ
- Sample intermediate state ***x***’**_t+1_** from application of ***T***_***t***_ to ***x***_***t***_
- Sample state ***x***_***t***+**1**_ after applying offspring formula.
- *t* ← *t* + *τ*
- **end while**

For clones where the time of barcoding is *s* > 0, we used Algorithm 1 to simulate a clone from time 0 up to time *s*, to obtain a distribution of cells ***x̃***_***s***_. From ***x̃***_***s***_, a cell was sampled at random ***x***_***s***_ = **1***_X_s__* where *X*_P_ ∼ ***x̃***_***s***_/|***x̃***_***s***_|_1_. Then starting from ***x***_***s***_, Algorithm 1 was used to simulate up to *t* = 1.

For each progenitor stage *s*, the output of the clonal dynamics model is a vector of counts of cells with a common progenitor at time *t* = 1, representing cells the time of measurement:

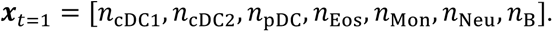

#### Model fitting

This model describes a probability distribution on the space 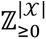 for each progenitor stage, from which samples can be accessed by sampling. Applying the log(1 + *x*) transform and clone-wise normalisation as described earlier, this becomes a probability distribution on **Δ**^|𝓧|^.

For progenitor stage *s*, let 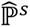 be the empirical distribution of fate profiles (i.e. an empirical distribution on the fate simplex), and let 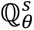 be the distribution of fate profiles induced by our model for some set of parameters *θ*. In order to fit our model, we aim to find parameters *θ* that minimise a data-fitting loss

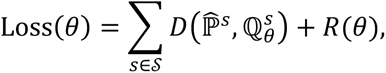

where *D*(*α*, *β*) is a divergence between probability distributions, and *R*(*θ*) is a regularisation term on the model parameters *θ*. In practice we do not have access to 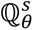 directly, but only via sampling, and so we fit based on 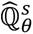, the empirical distribution constructed from repeated sampling from our model. In all, the loss function we seek to minimise in *θ* is

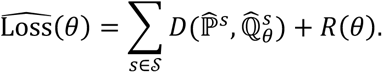

As both 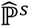 and 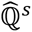 are empirical distributions, standard divergences such as Kullback-Leibler or Euclidean distances cannot be straightforwardly applied. We opt therefore to use the unbalanced Sinkhorn divergence (Séjourné et al., 2023) which by its nature handles the disjoint supports of its inputs. We use *λ* = 5 for the soft marginal constraint and *ε* = 0.05 for the entropic regularisation.

Model fitting is carried out by minimising 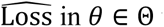 using covariance matrix adaptation evolution strategy (CMA-ES), a gradient-free optimisation algorithm that has shown good behaviour and robustness to noisy objective functions (Hansen et al., 2003).

The landscape in our model is parameterised by {Ψ*_ij_*, *i* ≥ *j*} and {Ψ*_i_*}, corresponding to the energy of transition states and attractors respectively, subject to the additional constraint that Ψ*_ij_* ≥ minWΨ*_i_*, Ψ*_j_*Y in order to ensure separation between any pair of attractors.

To parameterise the landscape in a way that incorporates the constraints on Ψ*_ij_*, we introduce variables *Z_ij_* ≥ 0, *i* ≤ *j* and *V_i_*, 1 ≤ *i* ≤ |𝒳| − 1. Then, Ψ*_ij_* = *Z_ij_* + maxWΨ*_i_*, Ψ*_j_*Y.

To parameterise the Ψ*_i_*, we observe that the dynamics induced by the potential landscape are invariant under the mapping Ψ*_i_* ↦ Ψ*_i_* + *C*, Ψ*_ij_* ↦ Ψ*_ij_* + *C*. Therefore we include the additional constraint that ∑*_i_* Ψ*_i_* = 0. To achieve this, the Ψ*_i_* are represented in terms of an orthogonal basis of the subspace ∑*_i_* Ψ*_i_* = 0, the coefficients of which are stored in *V_i_*.

The agent-based model requires that starting cells are sampled following a probability distribution ***p***_***0***_. While this could be taken as e.g. uniform, we choose to make our model more flexible by also allowing ***p***_***0***_ to be fit to data. We choose ***p***_***0***_ = softmax(***π***), where the entries are bounded −1 ≤ *π_i_* ≤ 1.

Division rates are an essential component of our model; since in our empirical data we only have cell counts, we implement only cell division in our agent-based model. We estimated cell proliferation rates independently for each cell type and each of the progenitor stages we consider, to capture (i) differences in proliferation rates between celltypes and (ii) proliferation of populations due to the irradiation and transplantation of progenitors. For a progenitor stage *s*, the average biomass produced for each celltype was calculated and log(1 + *x*)-transformed. To calculate an effective division rate for each celltype, this quantity was divided by the simulated timespan between the barcoding and readout, i.e. 1 − *s* celltype *i* at progenitor stage *s* is therefore 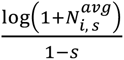 where 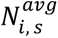 denotes the average biomass produced for celltype *i* for progenitor stage *s* .

Fitting a landscape to limited snapshot data is challenging, since there may in general be many landscapes that explain the data equally well. To counter this, we employed a regularisation term on the landscape parameters 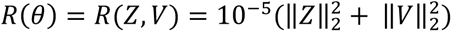.

Finally, to be robust to potential local minima in the loss function, we run CMA-ES with 256 random starting parameter configurations in the following ranges:

- 1 ≤ *Z_ij_* ≤ 10,
- −5 ≤ *V_i_* ≤ 5.

and for 512 iterations each to fit the model parameters (*Z*, *V*, *π*). We used a population size of 32. Candidate solutions were then ranked by their loss values averaged over the last 25 iterations.

---

**Supplementary Figure:**
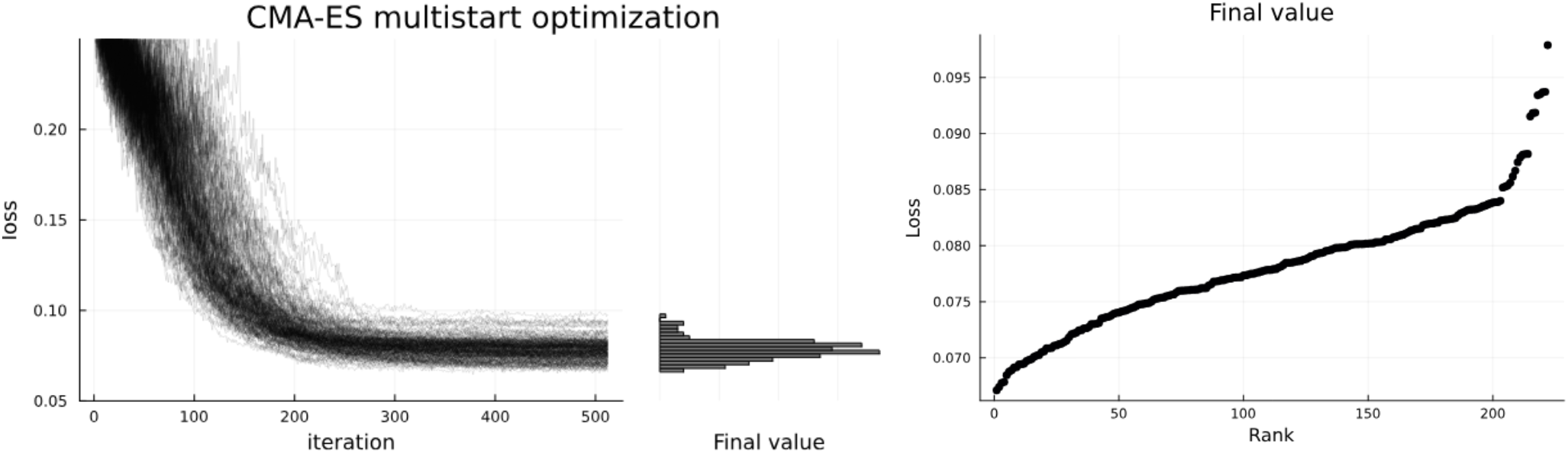
optimisation traces of each CMA-ES run, along with the distribution of final loss values (averaged over final 25 iterations of the algorithm), together with “waterfall” plot, in which candidates are ranked in terms of their final loss.

**Supplementary figure:**
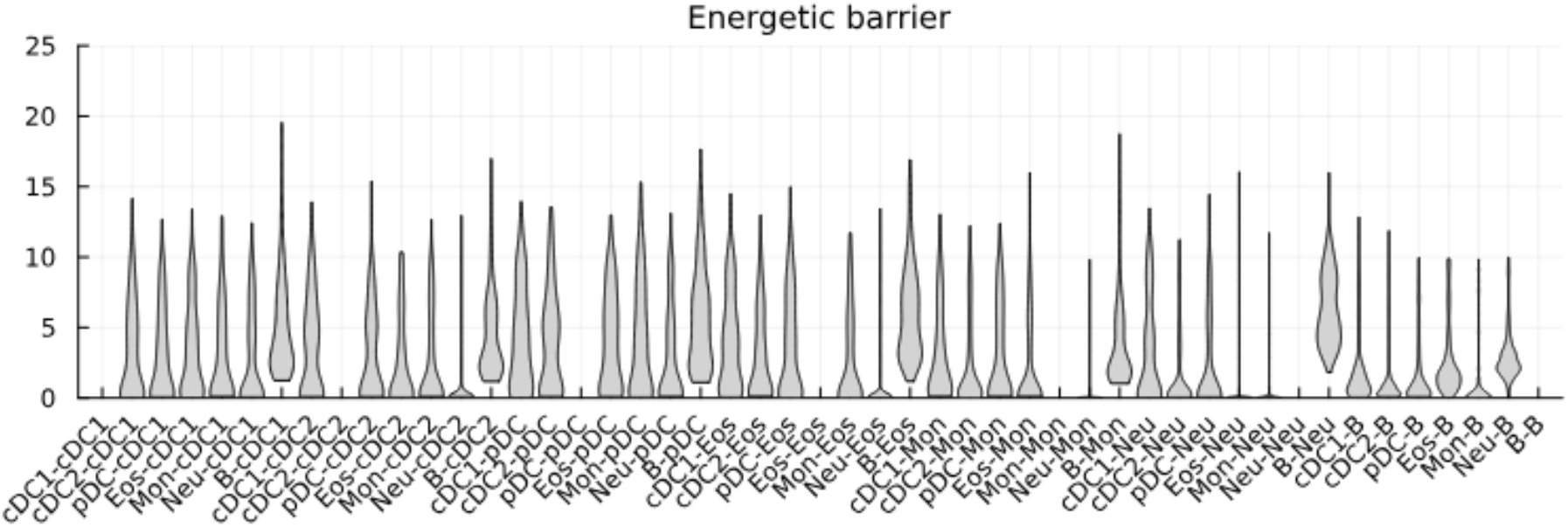
Distribution of energetic barriers between pairs of attractors in the landscape model, shown over all CMA-ES runs.

**Supplementary figure:**
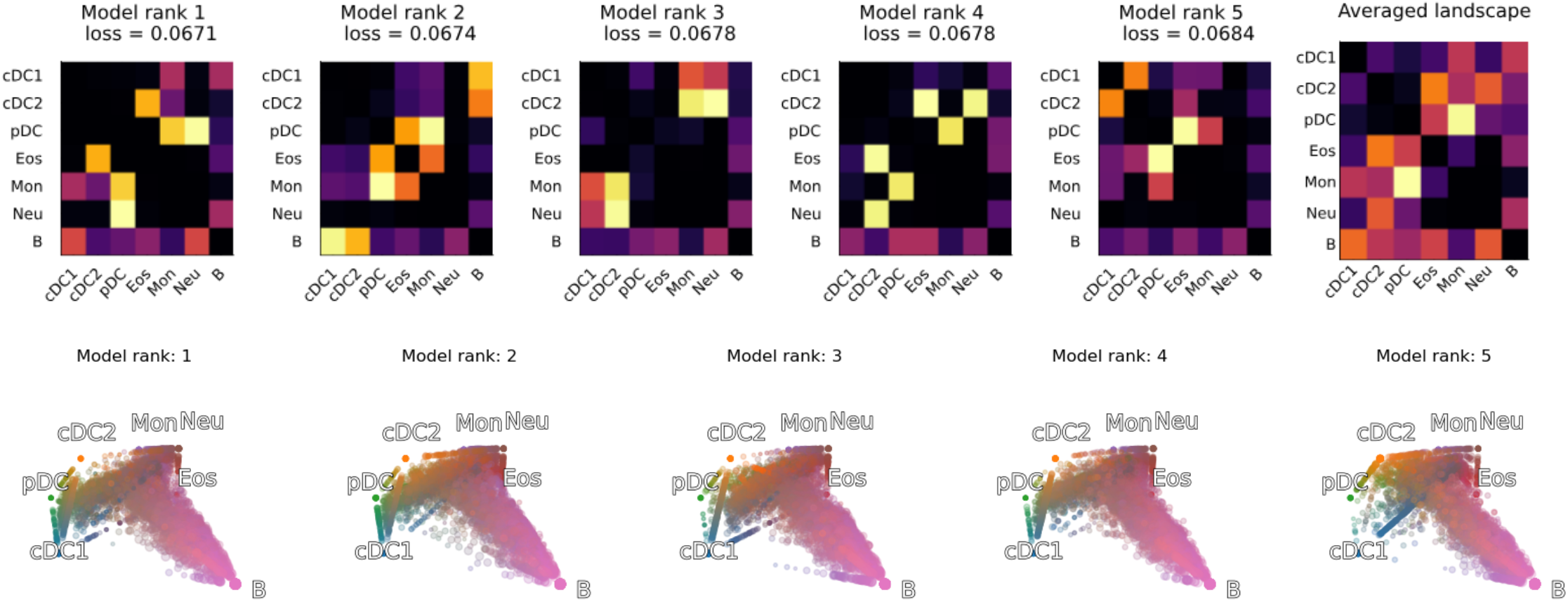
(a) Five best landscapes in terms of final loss (averaged over final 25 iterations of CMA-ES). (b) Average of five best landscapes. (c) Model outputs corresponding to five best landscapes.

## Data accessibility

Data and scripts to reproduce modelling results in Figures 5 and 6 are available at https://github.com/zsteve/multitrack_clonal_haematopoiesis.

## Acknowledgements

We thank the following WEHI facilities: Flow Cytometry, Genomics, Animal House. Also thanks to S.Tomei for technical assistance, and S.Tomei and S.Nutt for critical reading of the manuscript. This work was funded by National Health and Medical Research Council, Australia (1184736 (T.S.W. & S.H.N), 2009675 and 1145184 (S.H.N.)), and the Human Frontiers Special Program (RGP0060/2012).

## Contributions

J.S. performed all in vitro clonal fate (with J.T and A.N., and T.S. for analysis) and in vivo barcoding experiments. D.L. (with T.S.W.) performed filtering and visualization of the in vivo barcoding data, D.L. and S.H.N. interpreted the results, and S.Z. (with T.S.W) developed the mathematical model. S.H.N. and T.S.W. supervised the study. All authors contributed to writing of the manuscript.

---

**Figure S1.**
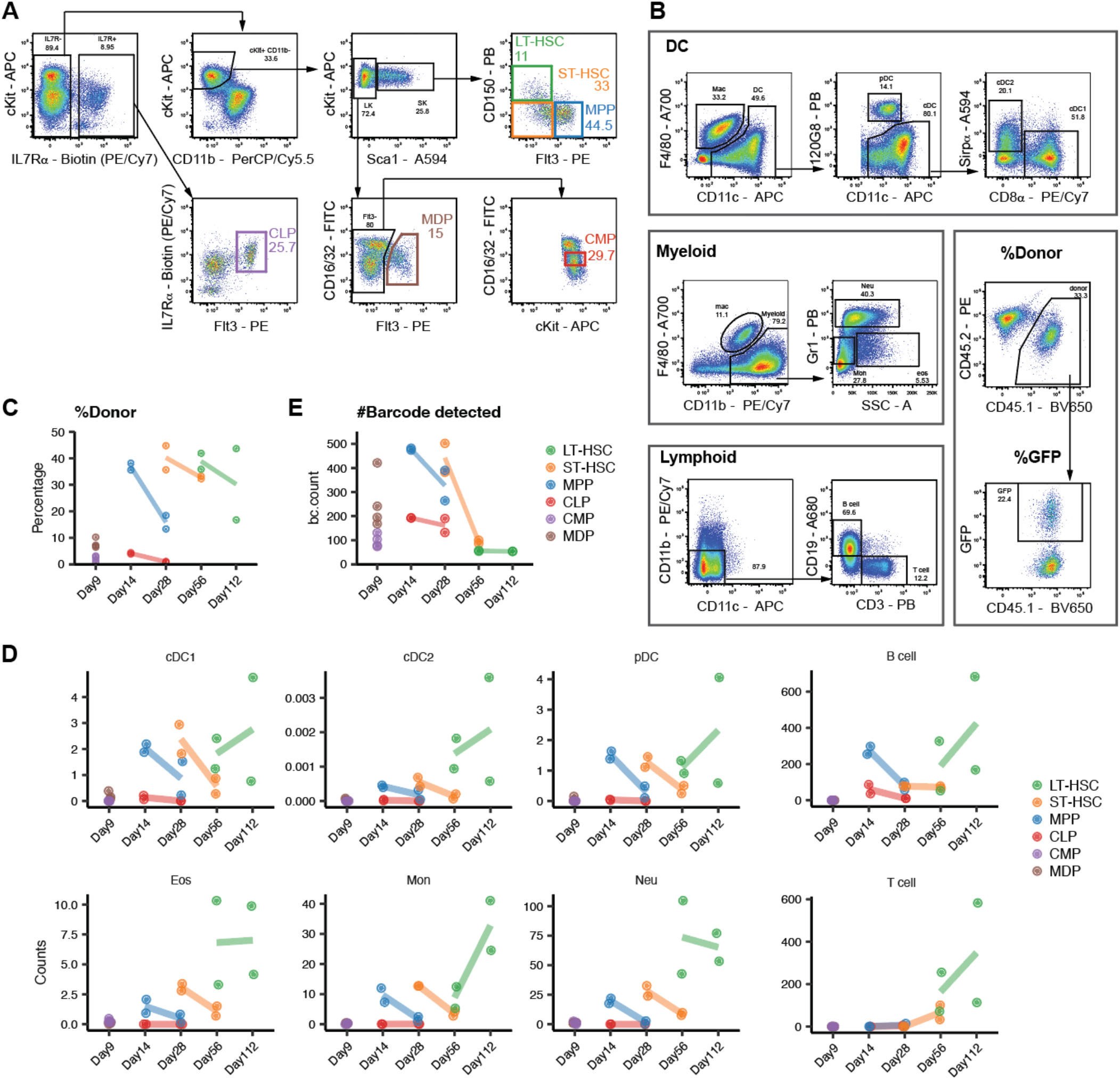
Gating strategy and barcoding metrics. A. Gating strategy to isolate different HSPC populations. B. Gating strategy to isolate different splenic progeny cell types. C. Percentage of donor engraftment per mouse. D. Fold expansion per cell type per mouse. E. Number of barcodes detected per mouse. (C-E) Each dot represents one recipient. Colour depicts HSPC group. Line connects the same HSPC group over different time point.

**Figure S2.**
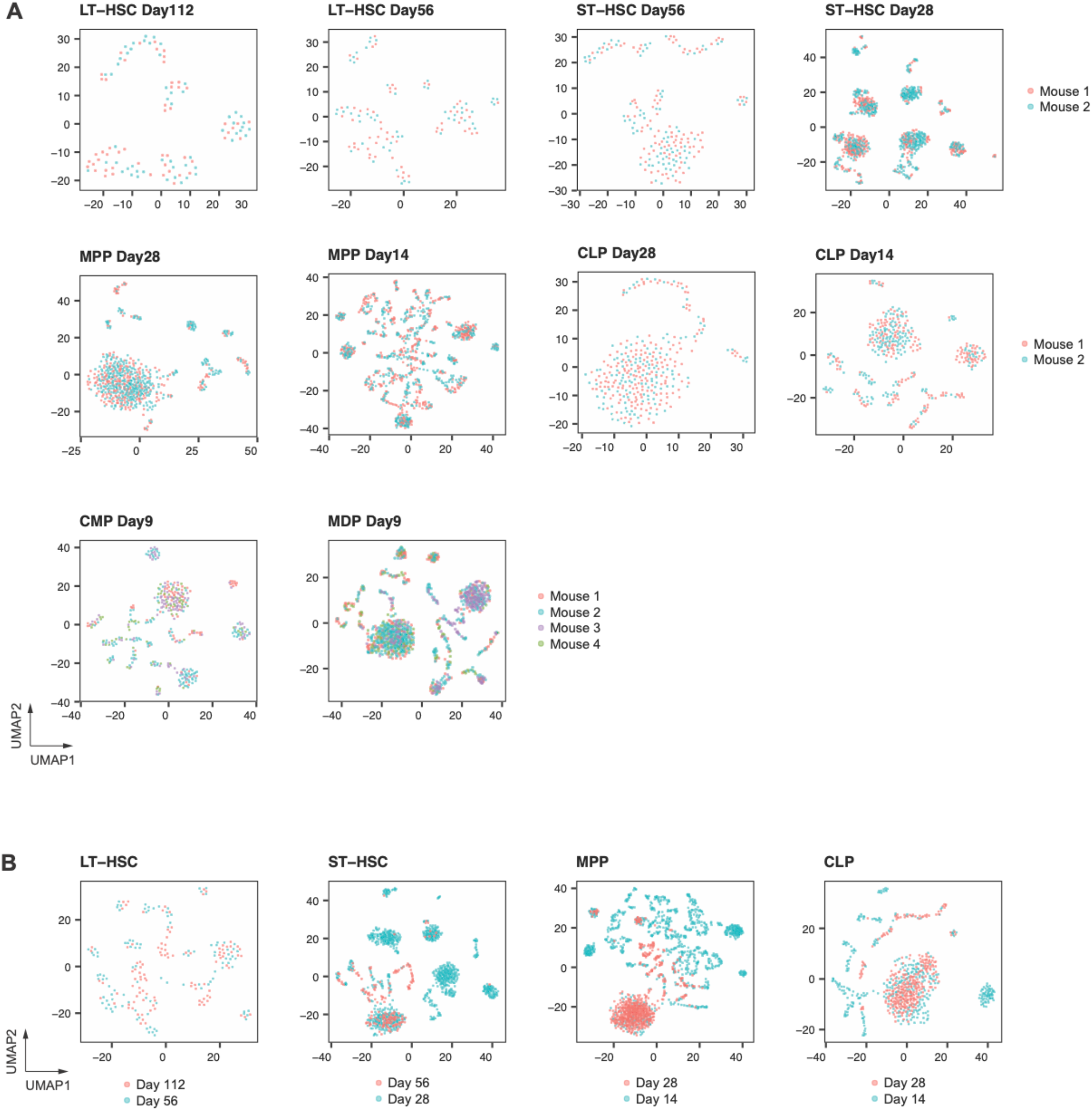
Biological reproducibility and time-related differences of HSPC clonal output. **A.** UMAPs of barcoding data for each HSPC subtype at each time point. Dot represents individual barcodes. Colour depicts biological replicate. **B.** UMAPs of barcoding data for each HSPC subtype pooled from different time points. Colour depicts different time points.

**Figure S3.**
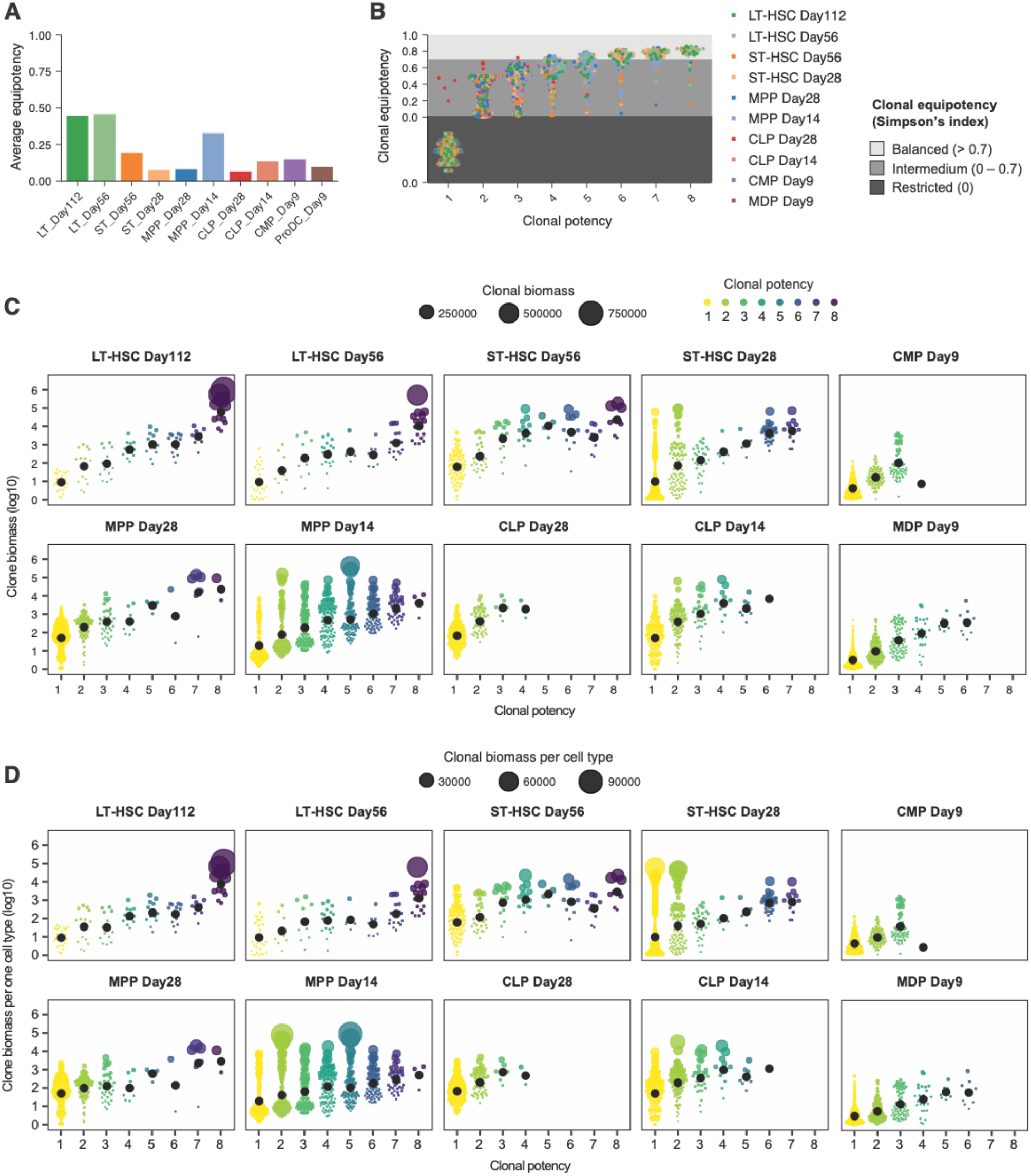
Heterogeneity in clonal features including potency, equipotency and biomass. A. Average equipotency (Simpson’s index) of barcodes from each HSPC/day group. B. Clonal equipotency of individual barcodes. Dot colour depicts HSPC/day group. C. Violin plots showing clonal biomass of individual barcodes (dots) with different clonal potency within each HSPC/day group. Dot size depicts clonal biomass (linear). D. Violin plots showing clonal biomass per cell type of individual barcodes (dots) with different clonal po-tency within each HSPC/day group. Dot size depicts clonal biomass per cell type (linear scale).

